# HARIBOSS++: An Integrated Platform for RNA-Targeted Drug Design

**DOI:** 10.64898/2026.09.07.749797

**Authors:** Matteo Marengo, Elodie Chapeaublanc, Hervé Ménager, Paraskevi Gkeka, Massimiliano Bonomi

## Abstract

Targeting RNA with small molecules is increasingly viewed as a pivotal strategy for future therapeutics. A major limitation in studying RNA-small molecule (RNA-SM) interactions is the lack of available experimental structures. HARIBOSS established a milestone as a curated database of RNA-SM complexes derived from the Protein Data Bank. Building on its community engagement, we present HARIBOSS++, an updated and next-generation platform designed to facilitate the study of future RNA-targeted drugs. Major updates include an extension of the pocketome to encompass binding sites where small molecules interact not only with RNA but also with other biomolecules, enhanced analysis of RNA molecules by family, detailed 3D structural characterization, improved analysis of ligands chemical space, addition of experimental binding affinity as well as molecular dynamics data, when available. HARIBOSS++ opens new avenues for biologists, bioinformaticians, and machine learning researchers to investigate RNA-SM interactions on a new scale. HARIBOSS++ can be explored using a web interface freely available at https://hariboss.pasteur.cloud.

## Introduction

RNA molecules perform a wide array of biological functions. Beside the well-known messenger RNA (mRNA), which conveys genetic information from DNA to proteins, several non-coding RNAs perform essential functions^1–3^. In fact, almost 70% of the genome codes for non-coding RNAs, while only 1.5% codes for proteins^4^. As a result, beyond the successful use of mRNAs in vaccines^5^, RNA offers a wide range of therapeutic opportunities through direct targeting of both coding and non-coding RNAs, thereby expanding the druggable space beyond traditional protein targets^4,6^. Historically, coding and non-coding RNAs have mainly been modulated using oligonucleotide-based approaches, including antisense oligonucleotides^7^, small interfering RNAs (siRNAs), and microRNAs (miRNAs)^8^.

In parallel, direct targeting of RNA with small molecules (SMs) has become an attractive alternative, as SMs often display better bioavailability, improved membrane penetration, and lower production costs^9,10^. Several RNA-targeting SMs have been developed, including Linezolid (Zyvox) in the early-mid 1990s^11^, which binds to ribosomal RNA to inhibit bacterial translation, and Ribocil from Merck in 2015^12^, which targets a flavin mononucleotide riboswitch to regulate gene expression at the translational level. A major milestone in the field is the approval by the Food and Drug Administration of Risdiplam in 2020, developed by Roche^13^, a small molecule that modulates SMN2 pre-mRNA splicing to treat Spinal Muscular Atrophy. The clinical pipeline is also expanding, with compounds in trials such as Zotatifin^14^, a translation controller that inhibits the translation of oncogenic mRNAs, or SKY-0515^15^, which targets Huntington disease. However, beyond these promising advances, some disappointments remain, such as the termination of the phase 2 clinical trial of Branaplam, developed by Novartis^16,17^. Despite these setbacks, the growing interest in RNA targeting has stimulated the development of new computational tools specific for RNA drug discovery.

Computer-aided drug design for RNA targets has progressed rapidly over the past few years. High-resolution structure prediction has recently advanced with deep learning based co-folding approaches such as Boltz2^18^ or AlphaFold3^19^, which model RNA-SM complexes directly from sequence. In parallel, structure-based virtual screening has benefited from docking and scoring strategies adapted to RNA. For instance, RNAmigos2^20^ proposes a graph learning framework for RNA virtual screening. RNA binding site detection has also improved with methods such as SHAMAN^21^, MVRBind^22^, GATRsite^23^, and DRLiPS^24^, among others^25^. More recently, RNA foundation models, including RNA-FM^26^, RiNALMO^27^, or ChaRNABERT^28^, have shown that useful information can be learned directly from RNA sequences^25,29–32^. These embeddings are now used by downstream methods such as DeepRNA-DTI^33^, RLSite^34^, and COBRA^35^ often in combination with graph-based representations to predict RNA binding sites. Despite these new methods, the development, benchmarking, and training of RNA-SM computational approaches remain constrained by the availability of curated and consistent structural datasets.

This lack of high-quality structural data originates from the limited number of experimentally determined RNA–SM structures available in the Protein Data Bank (PDB)^36^. Although the PDB contains close to 250,000 structures, most entries are protein-only (86%), whereas RNA-only structures (0.6%) and RNA-containing complexes (6%) represent a small fraction. This weakness motivated the creation of HARIBOSS^37^ (Harnessing Ribonucleic acid - Small Molecule Structures) as a curated database dedicated to RNA-SM interactions. Since its release in 2022, HARIBOSS has been widely used by the community, for example to train and evaluate binding-site prediction methods such as GerNA-Bind^38^, SmartBind^39^, E3NN^40^, RNABind^41^, or fpocketR^42^. Schrödinger also used it to evaluate and improve their tools for RNA (SiteMap^43^, Glide^44^, or AB-FEP+^45^). This large adoption indicates that enriching this database with additional information will further support emerging applications and a growing community. In addition, while previous works already analyze RNA-SM data^46,47^, or collect them in databases (NALDB^48^, SMMRNA^49^, RNALigands^50^, R-BIND^51,52^), they do not provide structural information as precise and curated as HARIBOSS^37^.

Here, we present an extension of HARIBOSS to better match current and future needs in RNA-focused drug discovery. In this new version of the database, we now include binding pockets in which the SM does not interact exclusively with RNA, in order to capture more diverse biological contexts. Furthermore, by enriching HARIBOSS with experimental binding affinity and molecular dynamics data, we aim to complement static structures with information on interaction strength and conformational variability. To further characterize binding pockets, we introduce a hierarchical classification of RNA families based on automated extraction of PDB metadata and provide improved analysis of RNA 2D structural diversity and ligand chemical space. These additions are now available in an updated HARIBOSS web interface, which is designed to provide clear entry points for biologists while also offering a curated, standardized database for bioinformatics and machine learning developments for RNA-targeted drug design.

## Methods

### HARIBOSS pipeline in a nutshell

The HARIBOSS pipeline consists of five successive steps (Fig. 1), described below and in detail in the original publication^37^.

1. **Fetching**: retrieve PDB entries containing RNA and at least one ligand.
2. **Filtering**: keep complexes containing drug-like compounds non-covalently bound to RNA.
3. **Clustering**: generate *redundant* and *non-redundant* versions of the extracted RNA-Small Molecule binding pockets.
4. **Analysis**: compute ligand and pocket descriptors using QikProp and SiteMap^53,54^.
5. **Visualization**: generate database statistics, plots, and update the HARIBOSS website.

**Figure 1:**
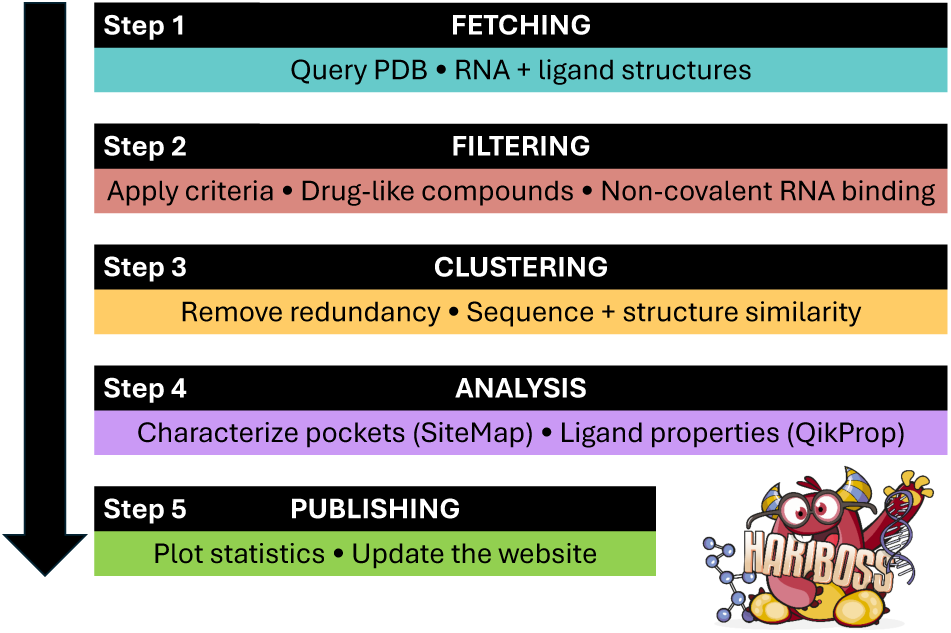
Overview of the HARIBOSS pipeline. The workflow is composed of five steps: RNA-small molecule complexes are first retrieved from the PDB (**fetching**, cyan), filtered according to predefined criteria (**filtering**, red), clustered into *redundant* and *non-redundant* versions (**clustering**, yellow), analyzed using Schrödinger tools (**analysis**, purple), and finally published on the HARIBOSS website (**publishing**, green).

### Expanding the Pocketome

To better represent the diverse composition of RNA binding pockets, we expanded the HARIBOSS beyond RNA-only environments. The ligand environment is defined as all residues within 6.0 Å of the ligand, including at least one RNA molecule (minimum 10 atoms), together with any proteins, DNA, ions, and other small molecules. Based on these interaction partners, pockets are assigned to one of 15 subcategories grouped into five major classes: RNA, RNA/Protein, RNA/DNA, RNA/Protein/DNA and RNA/Other. Here, biomolecule refers to any non-RNA molecular component of the environment - proteins, DNA, ions, or other small molecules - as detailed in Tab. S1. Some entries currently classified as RNA-only in fact correspond to RNA/RNA intermolecular pockets, most notably the splicing-modulator complexes 6HMO, 6HMI, 8R62, 8R63, 8R8P and 8CF2, in which the ligand bridges a pre-mRNA fragment and a U1 snRNA fragment. These structures form a distinct pocket class not reflected in our current five-way taxonomy and represent a natural addition for future HARIBOSS releases. mmCIF files are downloaded from the PDB and parsed with OpenMM (8.5.1)^55^ and MDAnalysis (v2.10.0)^56,57^, which classifies molecules as RNA, DNA, proteins, ions, water, or ligands. Modified residues are assigned to RNA, DNA, or protein according to their covalent connectivity. Drug-like ligands are kept if they satisfy the following criteria^58^: mass within 160 and 1000 Da; presence of at least one C atom; presence of only C, H, N, O, Br, Cl, F, P, Si, B, S and Se atoms. The Schrödinger structure Python API (v2026-2) saves two PDB files for each binding pocket, a local pocket including all residues within 12.0 Å of the ligand and a wider pocket taking the complete interacting chain(s). After this filtering stage, the database is clustered in a *non-redundant* version, as described in the next section.

### Hierarchical clustering of RNA-ligand binding pockets

To create a *non-redundant* version of the database, a hierarchical clustering approach was used. First we clustered based on RNA sequence and structural similarity. RNA interacting chains are clustered with CD-HIT-EST^59^ at 90% identity (chains shorter than 9 nucleotides are excluded). Sequences belonging to the same group, but interacting with different ligands, were considered as separate clusters. Within each group, we then clustered based on structural similarity of the pocket residues. To do this, a pairwise RMSD matrix over the backbone atoms of the pocket residues (sequence-aligned with ClustalW^60^) is computed and agglomerative clustering is performed with a 2.5 Å distance threshold cutoff. In presence of other biomolecules, like DNA or proteins, the same sequence-structure clustering procedure was recursively applied within each RNA cluster. After this hierarchical procedure, each cluster is represented by the structure with the highest experimental resolution.

### Binding pockets properties with Schrodinger suite

Binding-pocket geometry and physicochemical descriptors were computed with SiteMap^53,54^. This tool provides descriptors including SiteScore (ligandability), Dscore (druggability), hydrophilicity, hydrophobicity, enclosure, solvent exposure, volume and hydrogen-bond donor/acceptor character.

1. For each binding site, the corresponding receptor was prepared with the Schrödinger Protein Preparation Wizard (PrepWizard)^61^. Preparation was first attempted on the full receptor and, if unsuccessful or exceeding a predefined wall-time limit, repeated on the receptor truncated to residues within 12.0 Å of the ligand. Protonation states were assigned with Epik/PropKa^62^ at physiological pH (7.4), or at the experimental pH when available. Water molecules were removed, missing atoms and bond orders corrected, and hydrogen optimized with OPLS4 force field^63^ while heavy atoms remained restrained.
2. The prepared receptor and ligand were analyzed with SiteMap^53^ in the ligand-guided mode. SiteMap’s default compact mode is used; cavities larger than 800 Å^3^ were partitioned into compact sub-sites. When several candidate sites are returned, the representative site is the one that encloses most of the ligand, computed by Delaunay triangulation of the site-point cloud.
3. To improve robustness for the shallow, solvent-exposed cavities typical of RNA, a fallback strategy is applied: if the initial SiteMap calculation fails, it is automatically re-run with more permissive cavity-detection parameters; a reduced enclosure requirement (0.4 instead of 0.5) and a smaller van der Waals cutoff (0.55 instead of 1.1) that admits grid points closer to the receptor surface.
4. For nucleotide-dominated environments, the RNA-specific SiteMap scoring scheme was used^43^, rebalancing hydrophobic and hydrophilic contributions for RNA pockets and thus compensating for the intrinsically polar nature of RNA binding sites.

### Identification of RNA family classes

RNA structures were assigned to one of 12 biological families (Tab. S2) using a keyword-based classification pipeline (Fig. S1).

1. For each PDB entry, the corresponding structure webpage was queried using RCSB PDB API to retrieve macromolecule names, source organisms, chain identifiers, and entry titles.
2. Macromolecule names were matched against predefined keywords associated with each RNA family.
3. If no match was found, synthetic constructs were identified from the macromolecule name or source organism annotation. Otherwise, the PDB entry title was searched using the same keyword set, allowing biological annotations to override the synthetic-construct label when appropriate.
4. Chains matching none of these criteria were assigned to the *unknown* category.
5. Rfam^64^ annotations were then used as an independent chain-level cross-check only when the corresponding Rfam family could be mapped to one of the predefined RNA classes.
6. A pocket was assigned a single family when all annotated RNA chains belong to the same class, to *multiple* families when distinct classes were present, and labeled as *unknown* when no interacting RNA chain could be classified.

We acknowledge that this method could present some limitations (the PDB entry title keyword could potentially not correspond to all RNA chains of the complex, e.g splicing-modulator complexes that could contain both U1 snRNA and pre-mRNA). Its strength is that it gives to all chains a family type backed by keywords. Ideas to reinforce its quality could be to add some family types for a less coarse grained classification.

### RNA secondary-structure characterization

RNA secondary-structure types were annotated with x3DNA-DSSR^65^ (Dissecting the Spatial Structure of RNA). DSSR was run on each HARIBOSS structure, and its JSON output was parsed to assign each RNA nucleotide to one of these six structural categories: helix/paired, hairpin loop, internal loop, bulge, junction, or single-stranded. Annotations were also associated to pockets residues. For each structural context, enrichment within pockets relative to the complete RNA structure (only interacting chains were considered) was quantified as the log_2_ ratio of nucleotide coverage in pockets versus the whole RNA, where positive and negative values indicate enrichment and depletion, respectively. The statistical significance of each enrichment was assessed with a two-sided Fisher’s exact test and p-values were adjusted for multiple testing across the structure contexts using the Benjamini–Hochberg false discovery rate (FDR) procedure^66^.

### Exploration of chemical space of RNA binders

To provide an informative overview of the chemical space and ligand specificity, we used the TMAP^67^ visualization method. In this approach, molecules are encoded using molecular fingerprints (Min-Hash fingerprints (MHFP6)^68^ with a dimensionality of 2048 MinHash permutations) and indexed within a Locality-Sensitive Hashing (LSH) forest to enable efficient similarity search in high-dimensional space. Based on pairwise similarity relationships, an undirected weighted c-approximate k-nearest neighbor graph (c-k-NNG; k =20) is constructed, from which a minimum spanning tree is derived and embedded into two dimensions for visualization.

### Analysis of ligand properties

Ligands were evaluated against established drug-likeness criteria, including Lipinski’s Rule of Five^69^, Veber’s Rule^70^, and Pfizer’s 3/75 guidelines^71^ (Tab. S3). Two-dimensional molecular descriptors (MW, cLogP, HBD, HBA, TPSA, rotatable bonds, these abbreviations among others are defined in Tab. S4) needed to assess these rules were computed with RDKit^72^ from the canonical SMILES of each ligand. Additional physicochemical descriptors relevant to absorption, distribution, and overall drug-likeness were computed using QikProp (Tab. S5) from the corresponding three-dimensional structures provided in Structure Data File (SDF) format. Protonation states were those deposited in the PDB Chemical Component Dictionary, no pH-dependent ionisation model was applied.

### Curation of ligands for druggability analyses

The following rules were applied to identify small molecules that, while passing our selection criteria, are not commonly considered as drugs. First, we removed compounds typically introduced during X-ray crystallography, cryo-EM, or NMR sample preparation to fold, stabilize, phase, or otherwise preserve the RNA; genuine biological cofactors (Tab. S6) were retained as *bona fide* ligands. Second, we flagged detergent-like molecules carrying long aliphatic tails (*≥* 10 contiguous acyclic *sp*^3^ carbons). Third, we removed polyols, defined as molecules bearing *≥* 3 *sp*^3^ C-OH carbons in a chain. To avoid discarding legitimate ligands such as antibiotics, this last rule was applied only to polyol-dominated molecules, i.e. those with *≤* 12 heavy atoms (glycerol-/erythritol-class) or in which the polyol fragment accounts for *≥* 70% of all heavy atoms. This resulted in 14 SMs considered as non-relevant (Tab. S7). The associated pockets are kept in HARI-BOSS to ensure a complete overview of RNA binding pockets. Some analysis presented in the Results section do not consider these pockets and have been flagged as such.

### Binding Affinity data

To retrieve experimental binding affinity data, we used the PDB-bind database^73,74^, which offers a curated collection of binding constants for biomolecular complexes deposited in the PDB as a resource for characterizing the mechanisms of molecular recognition. The data utilized in this study were retrieved from the publicly accessible version of PDBbind+ (PDBbind v2020.R1),ensuring that the results are reproducible and accessible to the broader scientific community.

### Molecular Dynamics data

We integrated MD trajectory data into the HARIBOSS database by linking structures to MD simulations deposited in the Molecular Dynamics Data Bank (MDDB)^75^, a European repository of biosimulation data. Structures from the *rna-lig* collection associated to a recent paper that analyzes RNA-SM MD trajectories^76^ were retrieved via the MDDB REST API and completed with broader RNA-ligand searches to allow for future updates.

### Identification of related structures via BGSU equivalence classes

All HARIBOSS entries correspond to ligand-bound (*holo*) structures. To facilitate the identification of structurally equivalent *apo* and *holo* RNAs, we annotated each HARIBOSS entry with RNA structural equivalence classes from the Bowling Green State University (BGSU) RNA 3D Hub^77^. The RNA 3D Hub release (v4.48) was queried programmatically, and the corresponding non-redundant (NR) membership list (*allresolution*) was downloaded. In it, integrated functional elements (IFEs) are grouped into equivalence classes based on sequence and structural similarity. Each HARIBOSS entry was mapped to its equivalence class(es) and associated RNA chain(s), after which only the classes corresponding to RNA chains interacting with the ligand were kept.

### HARIBOSS website

The HARIBOSS website (https://hariboss.pasteur.cloud/), has undergone several updates. The compound-centric perspective is still available to identify compounds reported as RNA binders in HARIBOSS. On each entry page, one can find Canonical Smiles, IUPAC name, inChi, inchikey and SMARTS for each ligand. The three chemistry rules are reported along with a link to both Pubchem and Chembl. Ligands with five-letter identifiers are now correctly included. In addition, users can now filter structures by SMILES and SMARTS. In the complex-centric perspective, more substantial updates were implemented. For each pocket, the interaction environment is now reported. Interacting RNA chains are also annotated with their RNA family classification. In addition, when available, dissociation constants are provided in the pocket panel while molecular dynamics trajectories and related BGSU equivalence classes are provided in the complex panel. For all these new features, associated filters have been implemented. An interactive dashboard is provided to help users easily retrieve predefined filtered data on the Ligand and RNA-SM complexes pages. Finally, updated statistics reflecting the current state of the database are provided, including an interactive HTML TMAP representation of the ligands.

## Results

### Expansion of the RNA-SM pocketome and structural diversity

Based on June 19, 2026 update, we characterized HARIBOSS according to the composition of the pocket environment: complexes where ligands interact exclusively with RNA and complexes where it interacts with RNA as well as with other biomolecules. Our analysis shows that expanding the pocketome beyond RNA-only interactions significantly enriches the database (Tab. 1). In the redundant dataset, this expansion leads to an 82% increase in RNA-SM complexes, a 134% increase in identified pockets, and a 48% increase in unique ligands. Overall 43% of pockets involve RNA only, 38% RNA/Protein environment, 14% RNA/Other interactions, and the remaining involve DNA (Fig. 2A, Fig. S2). Heterogeneous interaction environments are common. For example, in PDB entry 8VRL^78^, chloramphenicol (CLM) simultaneously contains in its 6.0 Å binding pocket both RNA (10 nucleotides) and protein (7 residues), illustrating a typical RNA/Protein binding pocket (Fig. 2B). Additional examples of these diverse interaction environments are provided in Supplementary Figs (Fig. S3, Fig. S4 and Fig. S5).

**Figure 2:**
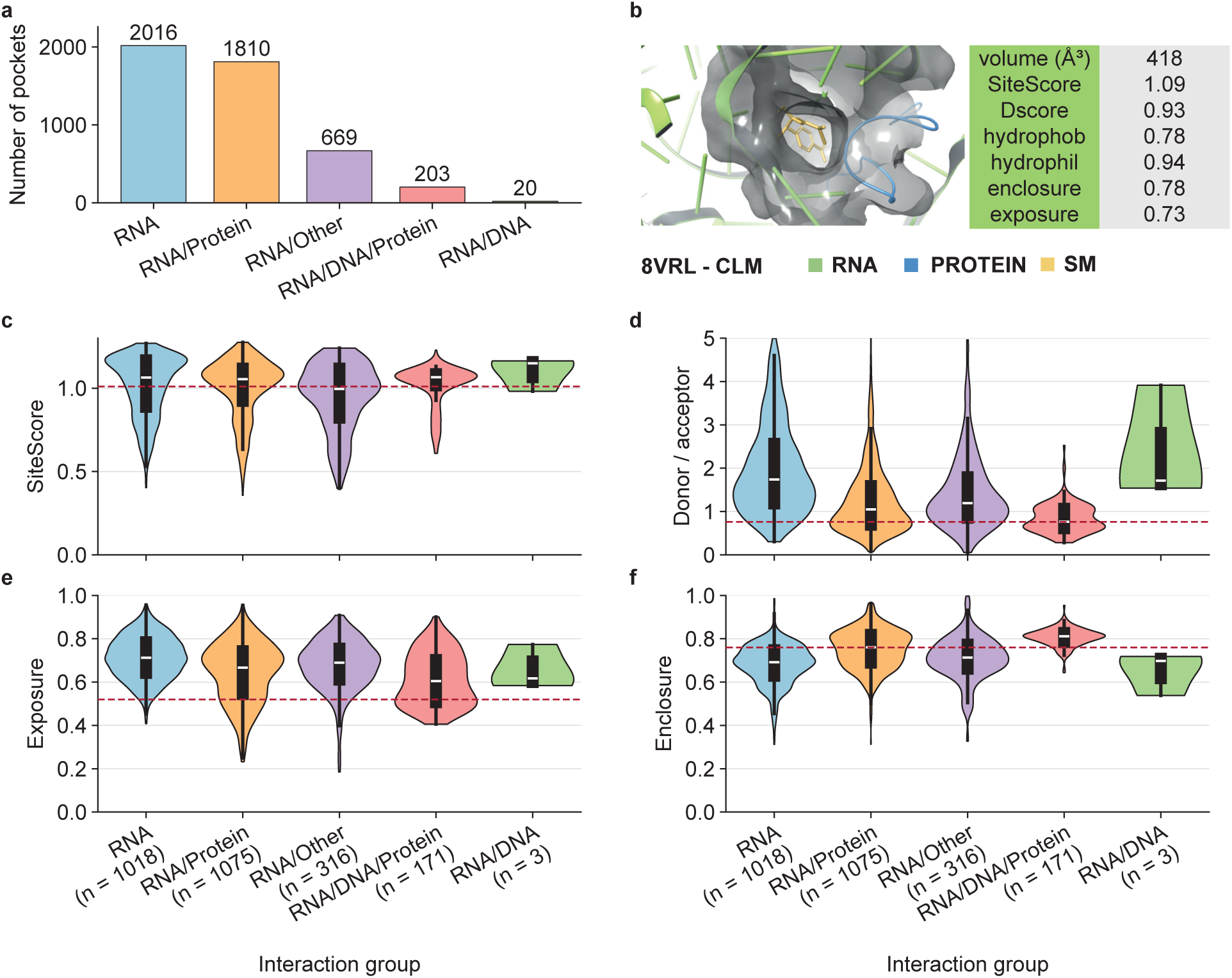
Binding pocket properties across interaction environments. **A.** Distribution of all binding pockets according to their interaction environment in the *redundant* HARIBOSS (*n* = 4718): RNA (light blue, *n* = 2016), RNA/protein (light orange, *n* = 1810), RNA/other (light purple, *n* = 669), RNA/DNA/Protein (light red, *n* = 203), RNA/DNA (light green, *n* = 20). **B.** Representative RNA/Protein binding pocket from PDB entry 8VRL^78^ where Chloramphenicol (CLM) binds at an interface of RNA and protein residues. **C & D.** Distribution of SiteScore and hydrogen bond donor/acceptor character across interaction environments. Dashed lines indicate references values computed for protein binding pockets^54^. **E & F.** Distribution of solvent exposure and pocket enclosure across interaction environments.

**Table 1:** Composition statistics of the HARIBOSS database. Reported value for the numbers of RNA-SM complexes, pockets and ligands. Data are categorized by structural composition of the binding pocket (RNA-only versus multi-biomolecule interfaces) and dataset redundancy.

| Structural Category | Dataset Type | RNA-SM complexes | Pockets | Ligands |
| --- | --- | --- | --- | --- |
| **RNA-only** | Redundant | 1114 | 2016 | 359 |
|  | Non-redundant | 883 | 1551 | 351 |
| **RNA (with or w/o partner)** | Redundant | 2025 | 4718 | 531 |
|  | Non-redundant | 1759 | 3946 | 517 |

The distribution of RNA nucleotides count per pocket (Fig. S6) shows that some pockets contain as few as one RNA nucleotide. In such cases, the pocket may be more influenced by protein or DNA chains than by RNA, which should be taken into account when interpreting the results. Additional statistics are provided in Supplementary Information, including database numbers after removing pockets associated to curated ligands (Tab. S8), deposition dates and experimental methods of the complexes (Fig. S7), proportion of molecule types by complexes (Fig. S8), mass distribution of the complexes in the *redundant* dataset (Fig. S9), and evolution of mass and composition repartition across the different steps of the database construction (Fig. S10).

### Properties of RNA-SM binding pockets

SiteMap^53^ successfully characterized 4184 of the 4718 binding pockets (88.7%). Most pockets involve a single RNA chain (83.8%), whereas 15.6% span two RNA chains and only 0.6% involve three or more (Fig. S11). To enable balanced comparisons across interaction environments, we restricted the analyses to our curated non-redundant pockets (2583 pockets, Tab. S8). RNA-only (*n* = 1018) and RNA/Protein (*n* = 1075) pockets were selected because they represent the two largest populations (Tab. 2). SiteScore is comparable between the two interaction environments (median 1.054 for RNA/Protein vs 1.064 for RNA-only; Fig. 2C), indicating that RNA-only and RNA/Protein binding pockets are, on average, equally able to accommodate drug-like compounds. A clearer separation is seen in the donor/acceptor character, which is significantly higher for RNA-only pockets (median 1.740 vs 1.047; Fig. 2D), highlighting the more polar nature of RNA (requiring ligands capable of forming extensive hydrogen-bonding networks). RNA-only pockets are also more solvent-exposed (median 0.712 vs 0.667; Fig. 2E) and less enclosed (median 0.692 vs 0.760; Fig. 2F) than RNA/Protein pockets. Consistently, RNA/Protein pockets are larger (median 435 vs 290 Å^3^), which may be explained by the presence of positively charged protein residues that stabilize RNA structures and allow the formation of larger pockets or grooves (Fig. S12). They also show a higher Dscore (median 0.920 vs 0.876).

**Table 2:** RNA-only vs RNA/Protein pocket SiteMap statistics. Curated, non-redundant pockets with a SiteMap evaluation; *n* = 2093; RNA-only *n* = 1018, RNA/Protein *n* = 1075.

|  | Value | SiteScore | Dscore | exposure | enclosure | phobic | philic | don/acc |
| --- | --- | --- | --- | --- | --- | --- | --- | --- |
| RNA-only | mean | 1.014 | 0.825 | 0.712 | 0.682 | 0.316 | 1.108 | 1.984 |
| RNA/Protein | mean | 1.001 | 0.854 | 0.643 | 0.751 | 0.295 | 1.286 | 1.252 |
| RNA-only | median | 1.064 | 0.876 | 0.712 | 0.692 | 0.196 | 1.049 | 1.740 |
| RNA/Protein | median | 1.054 | 0.920 | 0.667 | 0.760 | 0.214 | 1.255 | 1.047 |

This is backed up by the fact that RNA/Protein pockets are more often classified (Tab. S9) as difficult targets (0.83 *≤* Dscore *<* 0.98: 35.2% vs 25.7%), whereas RNA-only pockets fall more frequently into the undruggable category (Dscore *<* 0.83: 43.7% vs 34.3%). There are distinct advantages in targeting these ribonucleoprotein (RNP) complexes^79^, such as improved specificity and more stable pockets. Examples are the case of rocaglamide A (RocA) or marine macrolid pateamine A derivative (DMPatA) that binds at the interface of a complex between EIF4A1 and its RNA substrate (PDB 5ZC9^80^, PDB 6XKI^81^). Across all 2583 selected pockets, SiteMap descriptors show expected relationships (Fig. S13), including a negative correlation between druggability and hydrophilicity and a positive correlation between druggability and pocket volume. Complete SiteMap statistics for all interaction environments are provided in Supplementary Tab. S10.

### Analysis of RNA families

The RNA family annotation pipeline successfully identified a biological family for 4406 out of 4718 pockets (93.4%). Representative examples of HARIBOSS entries belonging to each RNA family are illustrated in Tab. S11. The distribution of RNA families across pockets (Fig. 3A, Tab. S12) shows that 60.9% of pockets are associated with rRNA, followed by riboswitches (9.4%), synthetic constructs (7.1%), aptamers (3.7%), tRNA (3.3%), and ribozymes (3.3%); only 6.6% of pockets remain classified as unknown. The strong predominance of rRNA-associated pockets likely reflects the experimental bias toward ribosomal structures in the PDB. In addition, 107 pockets involve multiple RNA families. PDB entry 5AA0^82^ illustrates such a case, where ligand 8AN binds at the interface between rRNA and tRNA (Fig. 3B). The ligand binds at the rRNA-tRNA interface because it captures the ribosome in the decoding step of translation. Such multi-RNA interfaces may represent underexplored binding environments and would support research for molecules targeting RNA-RNA interaction interfaces.

**Figure 3:**
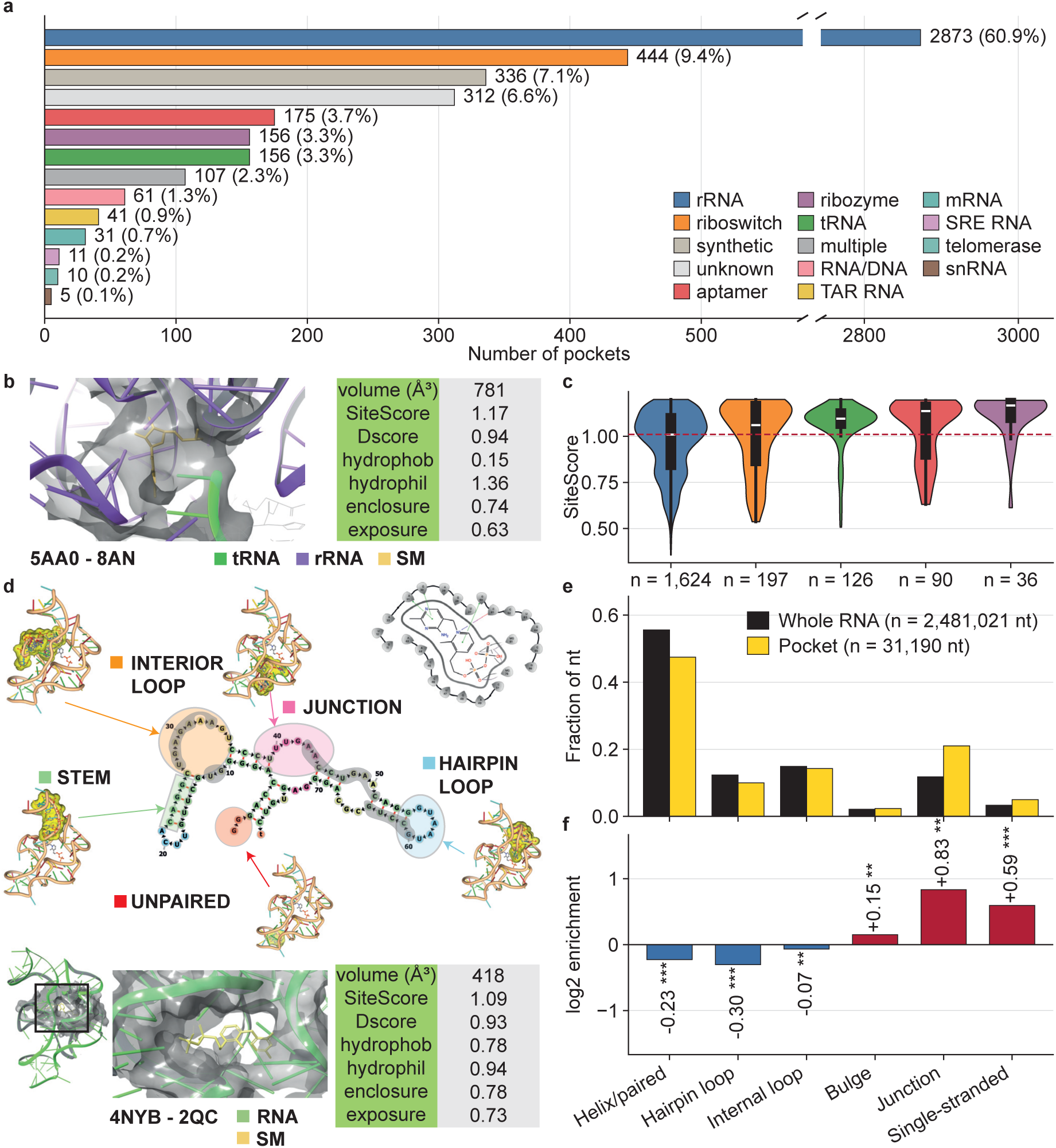
Analysis of RNA families and structural characterization of binding pockets. **A.** Distribution of RNA families across all *redundant* HARIBOSS binding pockets. **B.** Binding pocket from PDB entry 5AA0^82^ in complex with ligand 8AN. **C.** Distribution of SiteScore across rRNA (dark blue, *n* = 1624), riboswitch (orange, *n* = 197), tRNA (green, *n* = 126), aptamer (red, *n* = 90), ribozyme (purple, *n* = 36). **D.** Binding pocket in PDB entry 4NYB^83^ in complex with ligand 2QC along with its 2D interactions analyzed with DSSR^65^. **E.** Comparison of the fraction of nucleotides in each secondary-structure type between the entire interacting RNA chains and restricted to binding pockets. **F.** log2 enrichment between RNA secondary structure types in entire RNA chains vs binding pockets.

Analysis of SiteScore distributions across the five well-populated RNA families (curated pockets, *non-redundant*, *n* = 2073) shows that every family has a median SiteScore at or above 1.0, exceeding the canonical SiteMap threshold of 0.80 that separates drug-binding from non-drug-binding sites (Fig. 3C, Tab. S13). The highest median scores are found in RNA families that have evolved or been selected to bind small molecules: ribozyme (1.18), aptamer RNA (1.14) and riboswitch (1.12), followed by tRNA (1.10) and rRNA (1.03). This ordering could be biologically intuitive: riboswitches natively sense metabolites to regulate gene expression, ribozymes coordinate cofactors for catalytic activities, and in vitro-selected aptamers are optimised for shape-complementary ligand recognition, therefore all three tend to form well-defined pockets. The slightly lower (though still druggable) rRNA values reflect that many rRNA pockets are antibiotic binding sites embedded in a large, solvent-accessible ribosomal surface rather than purpose-built ligand cavities. In addition, within the RNA/Protein pockets most of them are rRNA (67 %) and most of these are ribosomal antibiotic sites that have been extensively studied. This selection effect has to be acknowledged so that if a model is trained on this data, DScore will inherit the bias. The remaining SiteMap descriptors are shown in Supplementary Fig. S14.

### Characterization of RNA secondary structure

Using DSSR^65^, we annotated the secondary-structure context of every RNA nucleotide, both across the entire RNA chains and within the binding pockets. PDB entry 4NYB^83^ with ligand 2QC illustrates that ligand-binding nucleotides occupy a restricted subset of the secondary-structure types present in the entire RNA structure (Fig. 3D). At the global scale, the RNA nucleotides occupy the full range of secondary structure types, from long helical stretches to hairpin loops, internal loops and junctions. The nucleotides in the ligand-binding pocket, however, occupy a more restricted subset, predominantly internal-loop, junction and adjacent paired positions.

We next compared the secondary-structure composition of RNA chains interacting with ligands to that of the corresponding binding pockets using the curated, *non-redundant* version of HARIBOSS (*n* = 2908 pockets, Fig. 3E&F). Canonical paired/helical nucleotides dominate both datasets but are less represented within pockets than across complete RNA chains (47.5% vs 55.6%; log_2_ enrichment = *-*0.23). In contrast, binding pockets are most strongly enriched in junctions (21.0% vs 11.8%; log_2_ = +0.83), followed by single-stranded nucleotides (5.0% vs 3.3%; log_2_ = +0.59) and, more modestly, bulges (2.3% vs 2.1%; log_2_ = +0.15). Conversely, hairpin loops are depleted within pockets (10.0% vs 12.3%; log_2_ = *-*0.30), and internal loops only marginally so (14.2% vs 14.9%; log_2_ = *-*0.07) (Tab. S14). Overall, RNA-SM binding pockets preferentially involves nucleotides located in flexible structural contexts, particularly single-stranded regions and junctions capable of forming cavities, consistent with previous analysis of RNA ligand-binding sites^84,85^.

### Small molecule specificity, chemical space, and druggability

A central question in targeting RNA is whether small molecules exhibit specificity toward a particular RNA families or interaction environments as selective ligands may serve as a therapeutic lead. We first investigated ligand specificity with respect to the interaction environment (Tab. S15, Fig. 4A). Among the 359 ligands observed in RNA-only interaction environments, 258 are exclusive to RNA, whereas 95 ligands are found exclusively in protein-associated environments. Similar trends are across RNA families (Tab. S16, Fig. 4B), with 162 of the 233 rRNA-associated ligands (70%) and 75 of the 101 riboswitch ligands (74%) found exclusively within their respective families.

**Figure 4:**
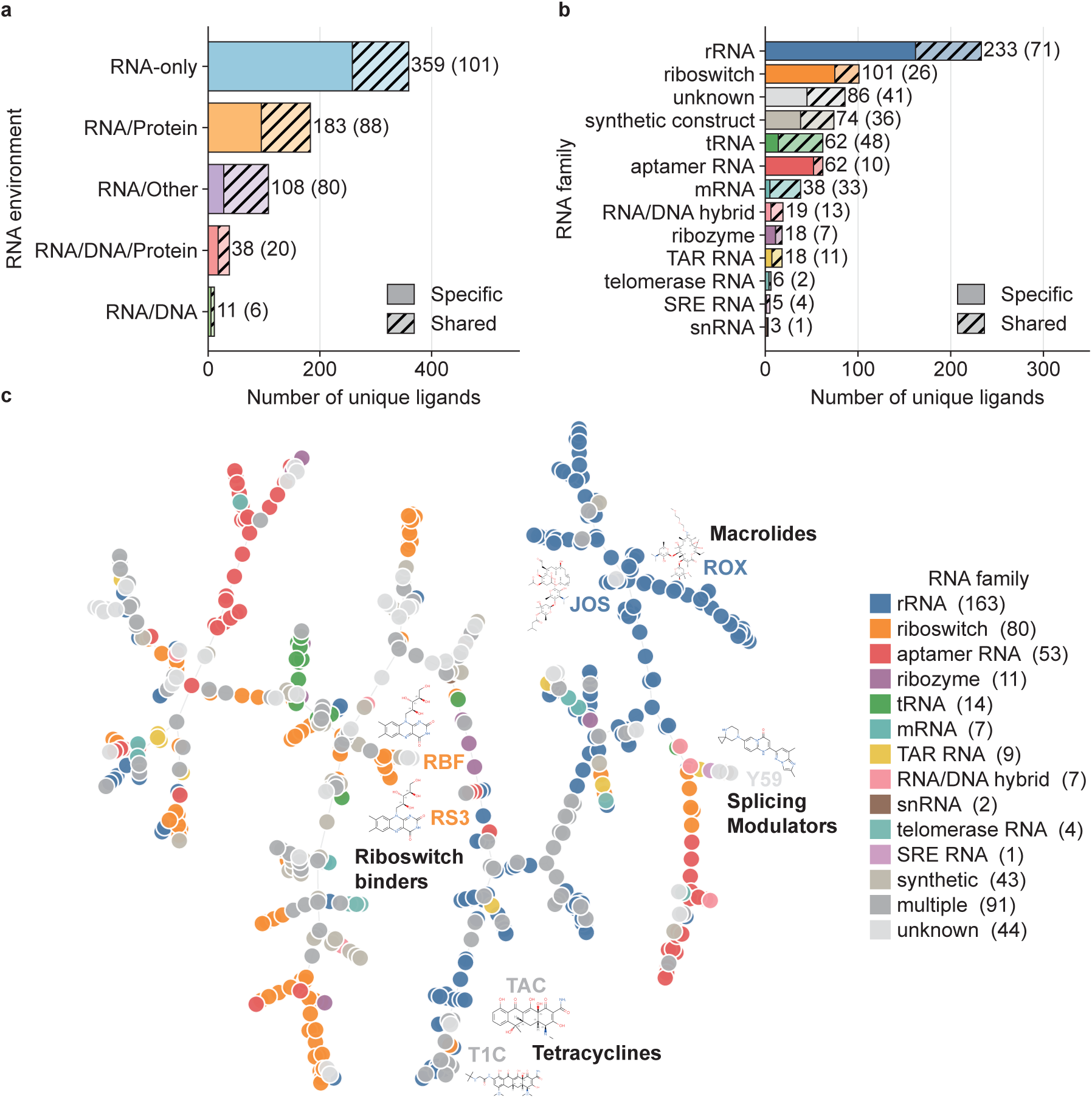
Characterization and visualization of the chemical space of RNA binders. **A.** Number of specific versus shared ligands for each RNA interaction environment. **B.** Number of specific versus shared ligands for each RNA family. **C.** TMAP visualization of the ligand chemical space (*n* = 529, 2 failed), colored according to the associated RNA family specificity.

To further explore these relationships, ligand chemical space was visualized using TMAP (Fig. 4C, Fig. S15). The resulting maps reveal distinct clusters associated with specific RNA families, suggesting that related RNA classes preferentially bind chemically similar compounds. Representative clusters include ribosome-targeting antibiotics (macrolides^86^, oxazolidinones, aminoglycosides and tetracyclines), riboswitch ligands (e.g., FMN in 1FMN, riboflavin (RBF) in 3F4G, roseoflavin (RS3) in 3F4H, and ribocil (6YG) in 5KX9), splicing modulators (Y59 and Y53), and amino acid-derived ligands (e.g. SAH and SAM). However, a subset of ligands, including linezolid (ZLD) and streptomycin (SRY), spans multiple RNA families, illustrating broader binding specificity.

Importantly, the interpretation of RNA ligand chemical space requires distinguishing pharmacologically relevant compounds from the curated list of commonly considered as non-drug-like molecules defined before. Spermine (SPM) is the most abundant member of this list, occurring in approximately 20% of all binding pockets, 68% of which belong to RNA/Protein environments (Tab. S17). Finally, drug-likeness analysis based on Lipinski’s and Veber’s rules shows that RNA-binding ligands are generally highly polar, with more than half exceeding the Veber topological polar surface area (TPSA, Fig. S16A) threshold. We also notice a low percentage of ligands having more than 10 rotatable bonds (Fig. S16C), highlighting long cavities that occur around the small molecules. Additional analyses include the ligand mass distribution (Fig. S17), the TMAP visualization by heterocycle content (Fig. S18) and QikProp physicochemical descriptors (Fig. S19).

### Annotating entries with binding affinity data

By interrogating the PDBbind+ database we identified 93 experimental binding affinity measurements associated with HARIBOSS entries, mainly dissociation constants (*K_d_*, n=87), but also 4 inhibition constants *K_i_*and 2 IC_50_ values. Most affinities (Fig. 5A) are grouped around 10^3^-10^4^ nM, indicating generally moderate binding affinities. The strongest interaction corresponds to ligand C2E bound to the bacterial c-di-GMP riboswitch (PDB entry 3MXH^87^;*K_d_* = 11 pM; Fig. 5E). In contrast, the weakest interaction is observed for arginine(ARG) bound to the HIV-1 TAR RNA 1ARJ^88^ (*K_d_*≈ 1 mM; Fig. 5F). This difference illustrates distinct RNA recognition modes. The interaction network between the RNA and the small molecule were characterized using the Protein Ligand Interaction Profiler (PLIP) ^89^ and the Schrodinger Interaction Profiler. In 3MXH, C2E occupies a riboswitch pocket and forms a dense interaction network, including hydrogen bonding, salt-bridge formation, and multiple π-stacking interactions with RNA bases. By contrast, arginine (ARG) in 1ARJ engages in limited electrostatic interactions, dominated by hydrogen bonds and π-cation contacts, and lacks any stacking interactions due to its non-aromatic nature (Fig. 5G).

**Figure 5:**
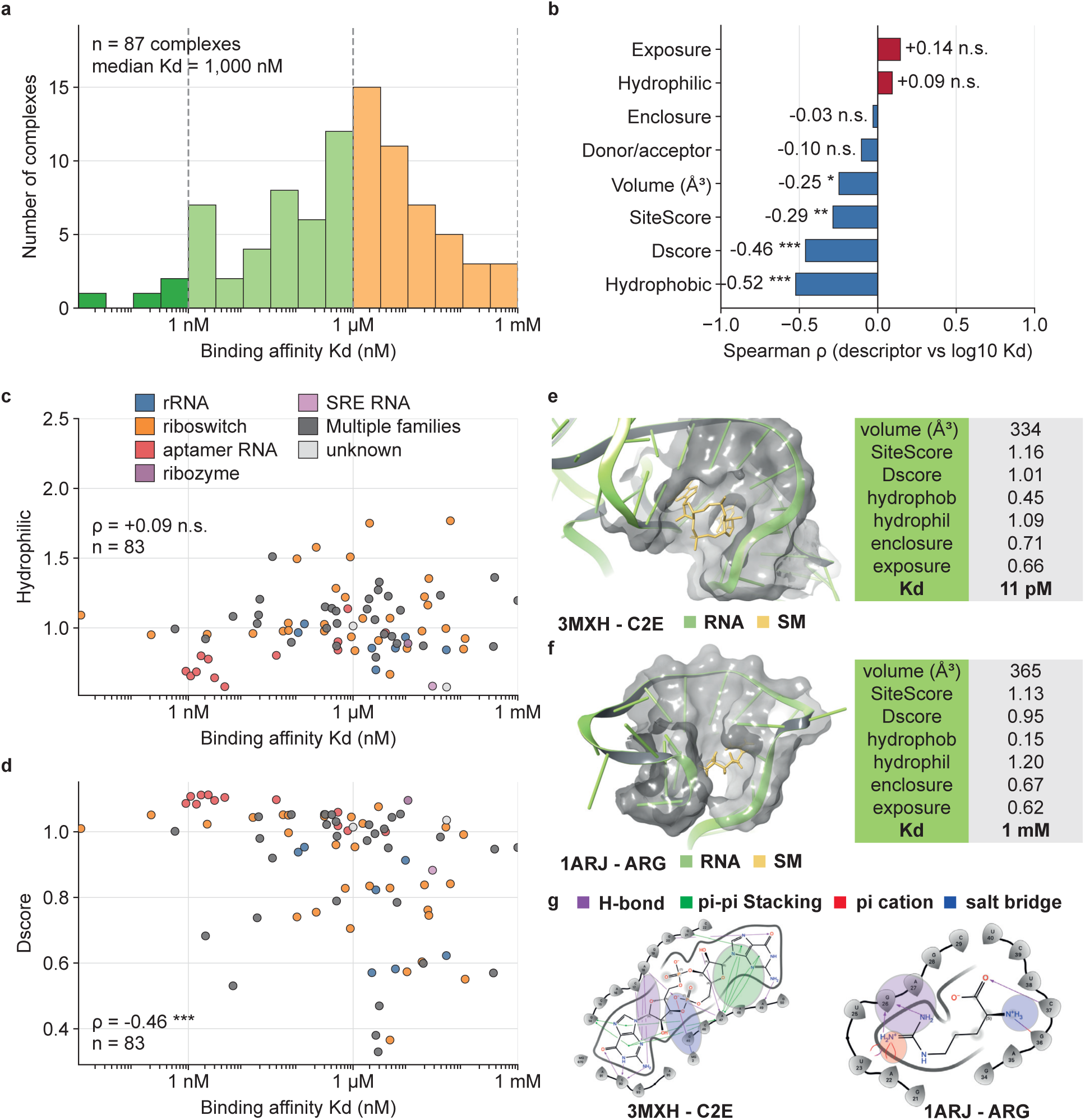
Integration of binding affinity data and analysis of correlations with physicochemical properties of ligands^53^. **A.** Distribution of experimental binding affinities for HARIBOSS ligands (*n* = 87). **B.** Spearman (rank) *ρ* of SiteMap metrics vs Binding Affinity correlation. **C.** Scatter plot of hydrophilicity versus binding affinity labeled by RNA families. **D.** Scatter plot of Dscore versus binding affinity labeled by RNA families. **E.** Detailed view of the binding pocket in PDB ID 3MXH^87^ in complex with c-di-GMP (C2E). **F.** Detailed view of the binding pocket in PDB ID 1ARJ^88^ in complex with arginine (ARG). **G.** Interaction analysis of the two binding pockets of PDB 3MXH and 1ARJ from Schrodinger Interaction profiler and PLIP^89^.

To investigate the relationship between pocket properties and binding affinity, Spearman (rank) correlations were computed between SiteMap descriptors (if multiple pockets within a complex, highest SiteScore one was selected as representative) and experimental dissociation constants. *K_d_* values were converted to nM and log-transformed (log_10_(*K_d_*)). The computed correlation (Fig. 5B, Fig. S20) reveals that binding affinity mostly correlates with pocket druggability and hydrophobic character. Dscore (*ρ* = *-*0.46) and hydrophobicity (*ρ* = *-*0.52) show the strongest negative correlations with log *K_d_*, indicating that more hydrophobic and druggable pockets are associated with stronger binding. SiteScore is also negatively correlated (*ρ* = *-*0.29), consistent with its integration of multiple pocket features. In contrast, hydrophilicity shows only a weak, non-significant positive trend with log *K_d_* (*ρ* = +0.09, n.s), providing a tentative indication that polar environments are less favorable for tight binding. The tightest binders (low *K_d_*) tend to occur at lower hydrophilic values, although the trend is modest (Fig. 5C). Geometric descriptors show weaker effects: larger pockets modestly favor binding (volume, *ρ* = *-*0.25), while solvent exposure is associated with slightly reduced affinity (exposure, *ρ* = +0.14, n.s), and enclosure has minimal impact (*ρ* = *-*0.03). Riboswitch-associated pockets tend to combine high Dscores with stronger binding affinities (Fig. 5D), although this observation should be interpreted cautiously given the limited number of experimentally characterized complexes (87 *K_d_*values but 83 reported as SiteMap failed for 4 pockets). The scatter plots with all SiteMap metrics are provided in Fig. S21.

Expanding the experimental affinity dataset is therefore essential to better characterize RNA-SM binding. While databases such as R-SIM^90^, SMMRNA^49^ or R-BIND^51,52^ provide valuable collections of RNA–SM binding data that can be used in computational pipelines (DeepRSMA^91^, RLAffinity^92^), they do not systematically link affinity measurements to experimental 3D structures. In this context, physics-based computational approaches such as AB-FEP+ (Schrodinger) appear promising, as recent studies report encouraging performance for estimating RNA–small molecule binding affinities^43^, whereas co-folding tools^18^ have not yet achieved the same accuracy. By progressively adding binding affinities to HARIBOSS, we aim to facilitate the development of structure-based approaches for the community and have a better understanding of the physicochemical interactions driving the recognition between small molecules and RNA.

### Integration of Molecular Dynamics data

Characterizing both structural and dynamic properties of RNA molecules is essential to identify selective compounds using structure-based approaches. However, accurately capturing RNA dynamics with molecular dynamics (MD) simulations remains challenging, in particular due to known limitations of current RNA force fields (recent benchmarks on the UUCG tetraloop and related systems highlight persistent limitations of RNA force fields in reproducing non-canonical pairs and hairpin geometry^93,94^). As of June 2026, trajectories are available for only eleven complexes^76^ (PDB IDs 1EI2, 2KGP, 1NTA, 1Q8N, 1UUI, 2L94, 5XI1, 6HMO, 7FHI, 6XB7, and 6VA3), generated using a variety of RNA force fields, including OL3^95^, OL3CP^93^, DES-AMBER and DESRES/DeShaw parameterizations^96^. These force fields were able to preserve the experimental RNA conformations, OL3cp demonstrating the best RNA secondary structure preservation. Concerning ligand binding, these force fields especially maintain stacking interactions but were suffering limited abilities for groove-binding ligands. Also to mention that large ribonucleoprotein assemblies (ribosomes, spliceosomal complexes) remain prohibitively expensive at atomistic resolution, restricting current MD coverage in HARIBOSS to smaller RNA–SM complexes.” Although currently limited in size, this integration represents an important step toward incorporating RNA dynamics in HARIBOSS, complementing static structural data. A detailed view of the different pockets classification around the different sets is presented through a Sankey diagram in Fig. S22.

### Apo–holo pairing of HARIBOSS complexes via RNA equivalence classes

Each of the HARIBOSS complexes was mapped to the *non-redundant*,*allresolution* equivalence classes defined by the BGSU RNA 3D Hub^77^. Considering every HARIBOSS entry as a holo structure, the other members of the corresponding equivalence class, not belonging to the database, can be considered as apo structures. This is a first, clean initial apo-holo mapping without any additional per-structure verification. Using this approach, an apo counterpart was found for 1377 holo structures (68.0%), while 493 (24.3%) belonged to a class containing no additional structure and 155 (7.7%) could not be assigned to any BGSU class. Representative examples include the FMN riboswitch complexes (PDB entries 3F30, 3F2Q, 3F2Y), which share the equivalence class NR all 47203.4 with the apo structure 6WJR, and the SAM-I riboswitch complexes (PDB entries 7EAF, 5FKF, 5FKH), which share the equivalence class NR all 84517.3 with the apo structure 3IQP. Therefore, mapping HARIBOSS onto BGSU equivalence classes and removing the holo members provides matched apo-holo structure sets for a large fraction of HARIBOSS complexes, offering a basis for future studies of RNA pockets.

## Conclusions

HARIBOSS is a comprehensive database of RNA-small molecule interactions, providing structural pocket information and ligand physico-chemical properties to support the development of binding site characterization and prediction tools. Indeed, HARIBOSS can be used as a curated train/test split for RNA-pocket foundation models and as positive/negative site sets for druggability classifiers. By incorporating heterogeneous RNA/protein/DNA binding environments, HARIBOSS can now capture a broader range of interaction contexts. RNA family annotations further allow stratified analyses of the data. Building on the secondary structure analysis presented here, future work could extend to deeper physico-chemical characterization and machine learning approaches to identify patterns underlying small-molecule binding. Integration of experimental binding affinities and molecular dynamics data will expand our understanding of RNA conformational ensembles. Analysis of the chemical ligand space also provides initial insights into ligand fingerprints associated with RNA specificity. RNA-SM data and its associated tools are constantly evolving to always better characterize these complexes, so is our database. As with any other RNA structure-based resource, HARIBOSS still reflects the limitations of the available data compared to proteins. In addition, it is not evenly distributed across RNA families and ligand chemotypes. An explicit caveat being for example that Dscore statistics are calibrated against a corpus dominated by rRNA and ribosomla-antiobiotic sites, so extrapolation to lncRNA, splicing, IRES or other underrepresented targets is out of distribution. Further developments will aim to always integrate more experimental data when available. Some explorative updates could be annotation of modified nucleotides when interacting with SM, a Rfam-based ^64^ family alignment, a less coarse-grained family classification to add some specificity and an explicit RNA/RNA-interface class (as flagged for the splicing modulators). HARIBOSS is openly accessible and will be continuously updated as new structures and supporting data become available. We hope it will serve as a useful resource for the RNA-small molecule research community.

## Acknowledgments

We acknowledge Floriane Puccio for her initial contribution to the RNA family classification pipeline. We also thank all the participants of the *Paris RNA Modeling Club* as well as the CECAM workshop *Decoding Allostery in Protein-Nucleic Acid Complexes: The AI Revolution* for providing valuable feedback on HARIBOSS, in particular Samuela Pasquali, Elisa Frezza, Élise Duboué-Dijon, Fabrice Leclerc, Guillaume Stirnemann, Modesto Orozco, and Giovanni Bussi. The Schrödinger team was also of precious help for adapting and using their tools to study RNA/ligand interactions, in particular David Papin, David Rinaldo, and Jõao Rodrigues. We would like to thank the IT Operations team of the IT department for its ongoing maintenance efforts, which ensure the Kubernetes cluster remains in perfect working order, facilitating the deployment of web applications hosted at Institut Pasteur. We also thank Bryan Brancotte for his invaluable help in deploying this application. Finally, we would like to thank all the researchers who found our database useful for training, testing, and validating novel computational methods for drug discovery.

## Competing Interests

M. Marengo and P. Gkeka are or were Sanofi employees and may own stocks in Sanofi. M. Bonomi declares no competing interests.

## Notes

### Competing Interest Statement

M. Marengo and P. Gkeka are or were Sanofi employees and may own stocks in Sanofi. The other authors declare no competing interests.

## References

(1) Cech, T. R.; Steitz, J. A. The Noncoding RNA Revolution—Trashing Old Rules to Forge New Ones. Cell 2014, 157, 77–94.

(2) Esteller, M. Non-coding RNAs in human disease. Nature Reviews Genetics 2011, 12, 861–874.

(3) Quinn, J. J.; Chang, H. Y. Unique features of long non-coding RNA biogenesis and function. Nature Reviews Genetics 2016, 17, 47–62.

(4) Warner, K. D.; Hajdin, C. E.; Weeks, K. M. Principles for targeting RNA with drug-like small molecules. Nature Reviews Drug Discovery 2018, 17, 547–558.

(5) Pardi, N.; Hogan, M. J.; Porter, F. W.; Weissman, D. mRNA vaccines — a new era in vaccinology. Nature Reviews Drug Discovery 2018, 17, 261–279.

(6) Overington, J. P.; Al-Lazikani, B.; Hopkins, A. L. How many drug targets are there? Nature Reviews Drug Discovery 2006, 5, 993–996.

(7) Neil, E. E.; Bisaccia, E. K. Nusinersen: A Novel Antisense Oligonucleotide for the Treatment of Spinal Muscular Atrophy. The Journal of Pediatric Pharmacology and Therapeutics 2019, 24, 194–203.

(8) Falese, J. P.; Donlic, A.; Hargrove, A. E. Targeting RNA with small molecules: from fundamental principles towards the clinic. Chemical Society Reviews 2021, 50, 2224– 2243.

(9) Chen, S.; Mao, Q.; Cheng, H.; Tai, W. RNA-Binding Small Molecules in Drug Discovery and Delivery: An Overview from Fundamentals. Journal of Medicinal Chemistry 2024, 67, 16002–16017.

(10) Gkeka, P.; Svensson, F.; Roca Magadán, C.; de Groot, M. J.; Jerome, S. V. Computational Hit Finding: An Industry Perspective. Journal of Medicinal Chemistry 2025, 68, 10507–10519.

(11) Hashemian, S. M. R.; Farhadi, T.; Ganjparvar, M. Linezolid: a review of its properties, function, and use in critical care. Drug Design, Development and Therapy 2018, 12, 1759–1767.

(12) Howe, J. A. et al. Selective small-molecule inhibition of an RNA structural element. Nature 2015, 526, 672–677.

(13) Ratni, H. et al. Discovery of Risdiplam, a Selective Survival of Motor Neuron-2 (SMN2) Gene Splicing Modifier for the Treatment of Spinal Muscular Atrophy (SMA). Journal of Medicinal Chemistry 2018, 61, 6501–6517.

(14) Rosen, E.; Sharma, M.; Berz, D.; Caswell-Jin, J. L.; Spira, A. I.; Fulgar, G. A.; Densel, M.; Rana, N.; Sperry, S.; Warner, D.; Meric-Bernstam, F. Phase 1/2 dose expansion study evaluating first-in-class eIF4A inhibitor zotatifin in patients with ER+ metastatic breast cancer. Journal of Clinical Oncology 2023, 41, 1080–1080, 2023 ASCO Annual Meeting abstract 1080.

(15) Skyhawk Therapeutics, Inc. A Phase 2/3 Randomized, Double-Blind, Placebo-Controlled Study to Evaluate the Efficacy and Safety of SKY-0515 in Participants With Huntington’s Disease (FALCON-HD). ClinicalTrials.gov Identifier: NCT07378644, 2026; Phase 2/3; recruiting as of 2026-08-04. Accessed: 2026-08-04.

(16) Borowsky, B. et al. Oral splicing modulator branaplam in Huntington’s disease: a phase 2 randomized controlled trial. Nature Medicine 2026, 32, 103–112.

(17) Krach, F.; Boerstler, T.; Neubert, S.; Krumm, L.; Regensburger, M.; Winkler, J.; Winner, B. RNA splicing modulator for Huntington’s disease treatment induces peripheral neuropathy. iScience 2025, 28, 112380.

(18) Passaro, S.; Corso, G.; Wohlwend, J.; Reveiz, M.; Thaler, S.; Somnath, V. R.; Getz, N.; Portnoi, T.; Roy, J.; Stark, H.; Kwabi-Addo, D.; Beaini, D.; Jaakkola, T.; Barzilay, R. Boltz-2: Towards Accurate and Efficient Binding Affinity Prediction. bioRxiv, 2025; Preprint.

(19) Abramson, J. et al. Accurate structure prediction of biomolecular interactions with AlphaFold 3. Nature 2024, 630, 493–500.

(20) Carvajal-Patiño, J. G.; Mallet, V.; Becerra, D.; Niño Vasquez, L. F.; Oliver, C.; Waldispühl, J. RNAmigos2: accelerated structure-based RNA virtual screening with deep graph learning. Nature Communications 2025, 16, 2799.

(21) Panei, F. P.; Gkeka, P.; Bonomi, M. Identifying small-molecules binding sites in RNA conformational ensembles with SHAMAN. Nature Communications 2024, 15, 5725.

(22) Chen, S.; Huang, Z.; Wang, Y.; Li, Y.; Tan, Y. S.; Deng, L.; Wu, M. MVRBind: multi-view learning for RNA-small molecule binding site prediction. Briefings in Bioinformatics 2025, 26, bbaf489.

(23) Sun, C.; Zhang, L.; Zhang, L.; Song, Y.; Ma, B.; Wang, Y. GATRsite: RNA–Ligand Binding Site Prediction Using Graph Attention Networks and Pretrained RNA Language Models. Journal of Chemical Information and Modeling 2025, 65, 8448–8461.

(24) Krishnan, S. R.; Roy, A.; Wong, L.; Gromiha, M. M. DRLiPS: a novel method for prediction of druggable RNA-small molecule binding pockets using machine learning. Nucleic Acids Research 2025, 53, gkaf239.

(25) Gao, J.; Zhuo, C.; Zeng, C.; Liu, H.; Zhao, Y. Advances in Methods for Accurate Prediction of RNA–Small Molecule Binding Sites: From Isolated to AI-Integrated Strategies. Pharmaceuticals 2025, 18, 1593.

(26) Chen, J.; Hu, Z.; Sun, S.; Tan, Q.; Wang, Y.; Yu, Q.; Zong, L.; Hong, L.; Xiao, J.; Shen, T.; King, I.; Li, Y. Interpretable RNA Foundation Model from Unannotated Data for Highly Accurate RNA Structure and Function Predictions. arXiv, 2022; Preprint.

(27) Penić, R. J.; Vla**s**ić, T.; Huber, R. G.; Wan, Y.; Sikić, M. RiNALMo: general-purpose RNA language models can generalize well on structure prediction tasks. Nature Communications 2025, 16, 5671.

(28) Morales-Pastor, A.; Vázquez-Reza, R.; Wieczór, M.; Valverde, C.; Gil-Sorribes, M.; Miquel-Oliver, B.; Ciudad, Á.; Molina, A. Character-level Tokenizations as Powerful Inductive Biases for RNA Foundational Models. arXiv, 2024; Preprint.

(29) Karroucha, W.; Oliver, C.; Stoven, V.; Mallet, V. Machine Learning for RNA-Targeting Drug Design. Journal of Chemical Information and Modeling 2026, 66, 7840–7861.

(30) Zhou, Y.; Chen, S.-J. Advances in machine-learning approaches to RNA-targeted drug design. Artificial Intelligence Chemistry 2024, 2, 100053.

(31) Wang, H.; Zhang, Y.; Chen, J.; Zhan, J.; Zhou, Y. Zero-shot benchmarking of RNA language models in structural, functional, and evolutionary learning. Briefings in Bioinformatics 2026, 27, bbag098.

(32) Sacco, G.; Bussi, G.; Sanguinetti, G. Machine learning for RNA secondary structure prediction: a review of current methods and challenges. RNA 2026, 32, 443–456.

(33) Bae, H.; Nam, H. DeepRNA-DTI: a deep learning approach for RNA-compound interaction prediction with binding site interpretability. Journal of Cheminformatics 2026, 18, 3.

(34) Sun, S.; Yang, J.; Gao, L.; Li, P.; Liu, Y. RNA language model and graph attention network for RNA and small molecule binding sites prediction. Bioinformatics 2025, 41, btaf447.

(35) Jang, W.; Shin, W.-H. CoBRA: compound binding site prediction using RNA language model. Briefings in Bioinformatics 2026, 27, bbaf713.

(36) Berman, H. M.; Westbrook, J.; Feng, Z.; Gilliland, G.; Bhat, T. N.; Weissig, H.; Shindyalov, I. N.; Bourne, P. E. The Protein Data Bank. Nucleic Acids Research 2000, 28, 235–242.

(37) Panei, F. P.; Torchet, R.; Ménager, H.; Gkeka, P.; Bonomi, M. HARIBOSS: a curated database of RNA-small molecules structures to aid rational drug design. Bioinformatics 2022, 38, 4185–4193.

(38) Xia, Y.; Li, J.; Chu, Y.-T.; Rao, J.; Chen, J.; Hua, C.; Yu, D.-J.; Chen, X.-C.; Zheng, S. Deciphering RNA–ligand binding specificity with GerNA-Bind. Nature Machine Intelligence 2025, 7, 1996–2008.

(39) Jiang, S.; Taghavi, A.; Wang, T.; Meyer, S. M.; Childs-Disney, J. L.; Li, C.; Disney, M. D.; Li, Y. Small Molecule Approach to RNA Targeting Binder Discovery (SMARTBind) Using Deep Learning Without Structural Input. bioRxiv, 2025; Preprint.

(40) Deng, Z.; Gu, R.; Bi, H.; Wang, X.; Zhang, Z.; Wen, H. Predicting Ligand–RNA Binding Using E(3)-Equivariant Network and Pretraining. Machine Learning in Structural Biology (MLSB) Workshop, 36th Conference on Neural Information Processing Systems (NeurIPS 2022). 2022.

(41) Zhu, W.; Ding, X.; Shen, H.-B.; Pan, X. Identifying RNA-small Molecule Binding Sites Using Geometric Deep Learning with Language Models. Journal of Molecular Biology 2025, 437, 169010.

(42) Veenbaas, S. D.; Felder, S.; Weeks, K. M. fpocketR: A Platform for Identification and Analysis of Ligand-Binding Pockets in RNA. ACS Chemical Biology 2026, 21, 151–159.

(43) Chopra, N.; Abramyan, A. M.; Si Chaib, Z.; Chen, W.; Xiong, Y.; Bochicchio, A.; Rinaldo, D.; Jerome, S. V.; Zhang, Y. Enabling In-Silico Hit Discovery Workflows Targeting RNA with Small Molecules. Journal of Chemical Information and Modeling 2025, 65, 7393–7398.

(44) Friesner, R. A.; Banks, J. L.; Murphy, R. B.; Halgren, T. A.; Klicic, J. J.; Mainz, D. T.; Repasky, M. P.; Knoll, E. H.; Shelley, M.; Perry, J. K.; Shaw, D. E.; Francis, P.; Shenkin, P. S. Glide: A New Approach for Rapid, Accurate Docking and Scoring. 1. Method and Assessment of Docking Accuracy. Journal of Medicinal Chemistry 2004, 47, 1739–1749.

(45) Chen, W.; Cui, D.; Jerome, S. V.; Michino, M.; Lenselink, E. B.; Huggins, D. J.; Beautrait, A.; Vendome, J.; Abel, R.; Friesner, R. A.; Wang, L. Enhancing Hit Discovery in Virtual Screening through Absolute Protein–Ligand Binding Free-Energy Calculations. Journal of Chemical Information and Modeling 2023, 63, 3171–3185.

(46) Hewitt, W. M.; Calabrese, D. R.; Schneekloth, J., John S. Evidence for ligandable sites in structured RNA throughout the Protein Data Bank. Bioorganic & Medicinal Chemistry 2019, 27, 2253–2260.

(47) Zhou, T.; Wang, H.; Zeng, C.; Zhao, Y. RPocket: an intuitive database of RNA pocket topology information with RNA-ligand data resources. BMC Bioinformatics 2021, 22, 428.

(48) Kumar Mishra, S.; Kumar, A. NALDB: nucleic acid ligand database for small molecules targeting nucleic acid. Database 2016, 2016, baw002.

(49) Mehta, A.; Sonam, S.; Gouri, I.; Loharch, S.; Sharma, D. K.; Parkesh, R. SMMRNA: a database of small molecule modulators of RNA. Nucleic Acids Research 2014, 42, D132–D141.

(50) Sun, S.; Yang, J.; Zhang, Z. RNALigands: a database and web server for RNA-ligand interactions. RNA 2022, 28, 115–122.

(51) Morgan, B. S.; Sanaba, B. G.; Donlic, A.; Karloff, D. B.; Forte, J. E.; Zhang, Y.; Hargrove, A. E. R-BIND: An Interactive Database for Exploring and Developing RNA-Targeted Chemical Probes. ACS Chemical Biology 2019, 14, 2691–2700.

(52) Donlic, A.; Swanson, E. G.; Chiu, L.-Y.; Wicks, S. L.; Juru, A. U.; Cai, Z.; Kassam, K.; Laudeman, C.; Sanaba, B. G.; Sugarman, A.; Han, E.; Tolbert, B. S.; Hargrove, A. E. R-BIND 2.0: An Updated Database of Bioactive RNA-Targeting Small Molecules and Associated RNA Secondary Structures. ACS Chemical Biology 2022, 17, 1556–1566.

(53) Halgren, T. A. Identifying and Characterizing Binding Sites and Assessing Druggability. Journal of Chemical Information and Modeling 2009, 49, 377–389.

(54) Halgren, T. New Method for Fast and Accurate Binding-site Identification and Analysis. Chemical Biology & Drug Design 2007, 69, 146–148.

(55) Eastman, P. et al. OpenMM 8: Molecular Dynamics Simulation with Machine Learning Potentials. The Journal of Physical Chemistry B 2024, 128, 109–116.

(56) Gowers, R. J.; Linke, M.; Barnoud, J.; Reddy, T. J. E.; Melo, M. N.; Seyler, S. L.; Domański, J.; Dotson, D. L.; Buchoux, S.; Kenney, I. M.; Beckstein, O. MDAnalysis: A Python Package for the Rapid Analysis of Molecular Dynamics Simulations. Proceedings of the 15th Python in Science Conference. Austin, TX, 2016; pp 98–105.

(57) Michaud-Agrawal, N.; Denning, E. J.; Woolf, T. B.; Beckstein, O. MDAnalysis: A toolkit for the analysis of molecular dynamics simulations. Journal of Computational Chemistry 2011, 32, 2319–2327.

(58) Stefaniak, F.; Bujnicki, J. M. AnnapuRNA: A scoring function for predicting RNA-small molecule binding poses. PLOS Computational Biology 2021, 17, e1008309.

(59) Li, W.; Godzik, A. Cd-hit: a fast program for clustering and comparing large sets of protein or nucleotide sequences. Bioinformatics 2006, 22, 1658–1659.

(60) Thompson, J. D.; Higgins, D. G.; Gibson, T. J. CLUSTAL W: improving the sensitivity of progressive multiple sequence alignment through sequence weighting, position-specific gap penalties and weight matrix choice. Nucleic Acids Research 1994, 22, 4673–4680.

(61) Sastry, G. M.; Adzhigirey, M.; Day, T.; Annabhimoju, R.; Sherman, W. Protein and ligand preparation: parameters, protocols, and influence on virtual screening enrichments. Journal of Computer-Aided Molecular Design 2013, 27, 221–234.

(62) Johnston, R. C.; Yao, K.; Kaplan, Z.; Chelliah, M.; Leswing, K.; Seekins, S.; Watts, S.; Calkins, D.; Chief Elk, J.; Jerome, S. V.; Repasky, M. P.; Shelley, J. C. Epik: pKa and Protonation State Prediction through Machine Learning. Journal of Chemical Theory and Computation 2023, 19, 2380–2388.

(63) Lu, C.; Wu, C.; Ghoreishi, D.; Chen, W.; Wang, L.; Damm, W.; Ross, G. A.; Dahlgren, M. K.; Russell, E.; Von Bargen, C. D.; Abel, R.; Friesner, R. A.; Harder, E. D. OPLS4: Improving Force Field Accuracy on Challenging Regimes of Chemical Space. Journal of Chemical Theory and Computation 2021, 17, 4291–4300.

(64) Ontiveros-Palacios, N.; Cooke, E.; Nawrocki, E.; Triebel, S.; Marz, M.; Rivas, E.; Griffiths-Jones, S.; Petrov, A.; Bateman, A.; Sweeney, B. Rfam 15: RNA families database in 2025. Nucleic Acids Research 2025, 53, D258–D267.

(65) Lu, X.-J.; Bussemaker, H. J.; Olson, W. K. DSSR: an integrated software tool for dissecting the spatial structure of RNA. Nucleic Acids Research 2015, 43, e142.

(66) Benjamini, Y.; Hochberg, Y. Controlling the False Discovery Rate: A Practical and Powerful Approach to Multiple Testing. Journal of the Royal Statistical Society: Series B (Methodological) 1995, 57, 289–300.

(67) Probst, D.; Reymond, J.-L. Visualization of very large high-dimensional data sets as minimum spanning trees. Journal of Cheminformatics 2020, 12, 12.

(68) Probst, D.; Reymond, J.-L. A probabilistic molecular fingerprint for big data settings. Journal of Cheminformatics 2018, 10, 66.

(69) Lipinski, C. A.; Lombardo, F.; Dominy, B. W.; Feeney, P. J. Experimental and computational approaches to estimate solubility and permeability in drug discovery and development settings. Advanced Drug Delivery Reviews 1997, 23, 3–25.

(70) Veber, D. F.; Johnson, S. R.; Cheng, H.-Y.; Smith, B. R.; Ward, K. W.; Kopple, K. D. Molecular Properties That Influence the Oral Bioavailability of Drug Candidates. Journal of Medicinal Chemistry 2002, 45, 2615–2623.

(71) Hughes, J. D. et al. Physiochemical drug properties associated with in vivo toxicological outcomes. Bioorganic & Medicinal Chemistry Letters 2008, 18, 4872–4875.

(72) Greg Landrum and other RDKit contributors RDKit: Open-source cheminformatics. 2026; Zenodo concept DOI (all versions).

(73) Liu, Z.; Su, M.; Han, L.; Liu, J.; Yang, Q.; Li, Y.; Wang, R. Forging the Basis for Developing Protein–Ligand Interaction Scoring Functions. Accounts of Chemical Research 2017, 50, 302–309.

(74) Wang, R.; Fang, X.; Lu, Y.; Yang, C.-Y.; Wang, S. The PDBbind Database: Methodologies and Updates. Journal of Medicinal Chemistry 2005, 48, 4111–4119.

(75) Amaro, R. E. et al. The need to implement FAIR principles in biomolecular simulations. Nature Methods 2025, 22, 641–645.

(76) Fernández Migens, P.; Serrano-Chacón, I.; Orozco, M.; Battistini, F. Success and Limitations of Current Force Fields for the Description of RNA–Ligand Complexes. The Journal of Physical Chemistry B 2025, 129, 11679–11692.

(77) Leontis, N. B.; Zirbel, C. L. In *RNA 3D Structure Analysis and Prediction*; Leontis, N. B., Westhof, E., Eds.; Nucleic Acids and Molecular Biology; Springer Berlin Heidelberg, 2012; Vol. 27; pp 281–298.

(78) Majumdar, S.; Kashyap, A.; Koripella, R. K.; Sharma, M. R.; Hurst-Hess, K.; Manjari, S. R.; Banavali, N. K.; Ghosh, P.; Agrawal, R. K. HflX-mediated drug resistance through ribosome splitting and rRNA disordering in mycobacteria. Proceedings of the National Academy of Sciences 2025, 122, e2419826122.

(79) Rhodes, C.; Balaratnam, S.; Yazdani, K.; Seshadri, S.; Schneekloth, J. S. Targeting RNA-protein interactions with small molecules: promise and therapeutic potential. Medicinal Chemistry Research 2024, 33, 2050–2065.

(80) Iwasaki, S. et al. The Translation Inhibitor Rocaglamide Targets a Bimolecular Cavity between eIF4A and Polypurine RNA. Molecular Cell 2019, 73, 738–748.e9.

(81) Naineni, S. K.; Liang, J.; Hull, K.; Cencic, R.; Zhu, M.; Northcote, P.; Teesdale-Spittle, P.; Romo, D.; Nagar, B.; Pelletier, J. Functional mimicry revealed by the crystal structure of an eIF4A:RNA complex bound to the interfacial inhibitor, desmethyl pateamine A. Cell Chemical Biology 2021, 28, 825–834.e6.

(82) Kumar, V.; Chen, Y.; Ero, R.; Ahmed, T.; Tan, J.; Li, Z.; Wong, A. S. W.; Bhushan, S.; Gao, Y.-G. Structure of BipA in GTP form bound to the ratcheted ribosome. Proceedings of the National Academy of Sciences 2015, 112, 10944–10949.

(83) Warner, K. D.; Homan, P.; Weeks, K. M.; Smith, A. G.; Abell, C.; Ferré-D’Amaré, A. R. Validating Fragment-Based Drug Discovery for Biological RNAs: Lead Fragments Bind and Remodel the TPP Riboswitch Specifically. Chemistry & Biology 2014, 21, 591–595.

(84) Veenbaas, S. D.; Koehn, J. T.; Irving, P. S.; Lama, N. N.; Weeks, K. M. Ligand-binding pockets in RNA and where to find them. Proceedings of the National Academy of Sciences 2025, 122, e2422346122.

(85) Guo, Z.-H.; Yuan, L.; Tan, Y.-L.; Zhang, B.-G.; Shi, Y.-Z. RNAStat: An Integrated Tool for Statistical Analysis of RNA 3D Structures. Frontiers in Bioinformatics 2021, 1, 809082.

(86) Nor Amdan, N. A.; Shahrulzamri, N. A.; Hashim, R.; Mohamad Jamil, N. Understanding the evolution of macrolides resistance: A mini review. Journal of Global Antimicrobial Resistance 2024, 38, 368–375.

(87) Smith, K. D.; Lipchock, S. V.; Livingston, A. L.; Shanahan, C. A.; Strobel, S. A. Structural and Biochemical Determinants of Ligand Binding by the c-di-GMP Riboswitch. Biochemistry 2010, 49, 7351–7359.

(88) Aboul-ela, F.; Karn, J.; Varani, G. The Structure of the Human Immunodeficiency Virus Type-1 TAR RNA Reveals Principles of RNA Recognition by Tat Protein. Journal of Molecular Biology 1995, 253, 313–332.

(89) Schake, P.; Bolz, S. N.; Linnemann, K.; Schroeder, M. PLIP 2025: introducing protein– protein interactions to the protein–ligand interaction profiler. Nucleic Acids Research 2025, 53, W463–W465.

(90) Krishnan, S. R.; Roy, A.; Gromiha, M. M. R-SIM: A Database of Binding Affinities for RNA-small Molecule Interactions. Journal of Molecular Biology 2023, 435, 167914.

(91) Huang, Z.; Wang, Y.; Chen, S.; Tan, Y. S.; Deng, L.; Wu, M. DeepRSMA: a cross-fusion-based deep learning method for RNA–small molecule binding affinity prediction. Bioinformatics 2024, 40, btae678.

(92) Sun, S.; Gao, L. Contrastive pre-training and 3D convolution neural network for RNA and small molecule binding affinity prediction. Bioinformatics 2024, 40, btae155.

(93) Mlýnský, V.; Kührová, P.; Pykal, M.; Krepl, M.; Stadlbauer, P.; Otyepka, M.; Banás, P.; Sponer, J. Can We Ever Develop an Ideal RNA Force Field? Lessons Learned from Simulations of the UUCG RNA Tetraloop and Other Systems. Journal of Chemical Theory and Computation 2025, 21, 4183–4202.

(94) Kührová, P.; Mlýnský, V.; Zgarbová, M.; Krepl, M.; Bussi, G.; Best, R. B.; Otyepka, M.; Sponer, J.; Banás, P. Improving the Performance of the Amber RNA Force Field by Tuning the Hydrogen-Bonding Interactions. Journal of Chemical Theory and Computation 2019, 15, 3288–3305.

(95) Zgarbová, M.; Otyepka, M.; Sponer, J.; Mládek, A.; Banás, P.; Cheatham, I., Thomas E.; Jurecka, P. Refinement of the Cornell et al. Nucleic Acids Force Field Based on Reference Quantum Chemical Calculations of Glycosidic Torsion Profiles. Journal of Chemical Theory and Computation 2011, 7, 2886–2902.

(96) Tan, D.; Piana, S.; Dirks, R. M.; Shaw, D. E. RNA force field with accuracy comparable to state-of-the-art protein force fields. Proceedings of the National Academy of Sciences 2018, 115, E1346–E1355.

