## Supplementary Information for "HARIBOSS++: An Integrated Platform for RNA-Targeted Drug Design"

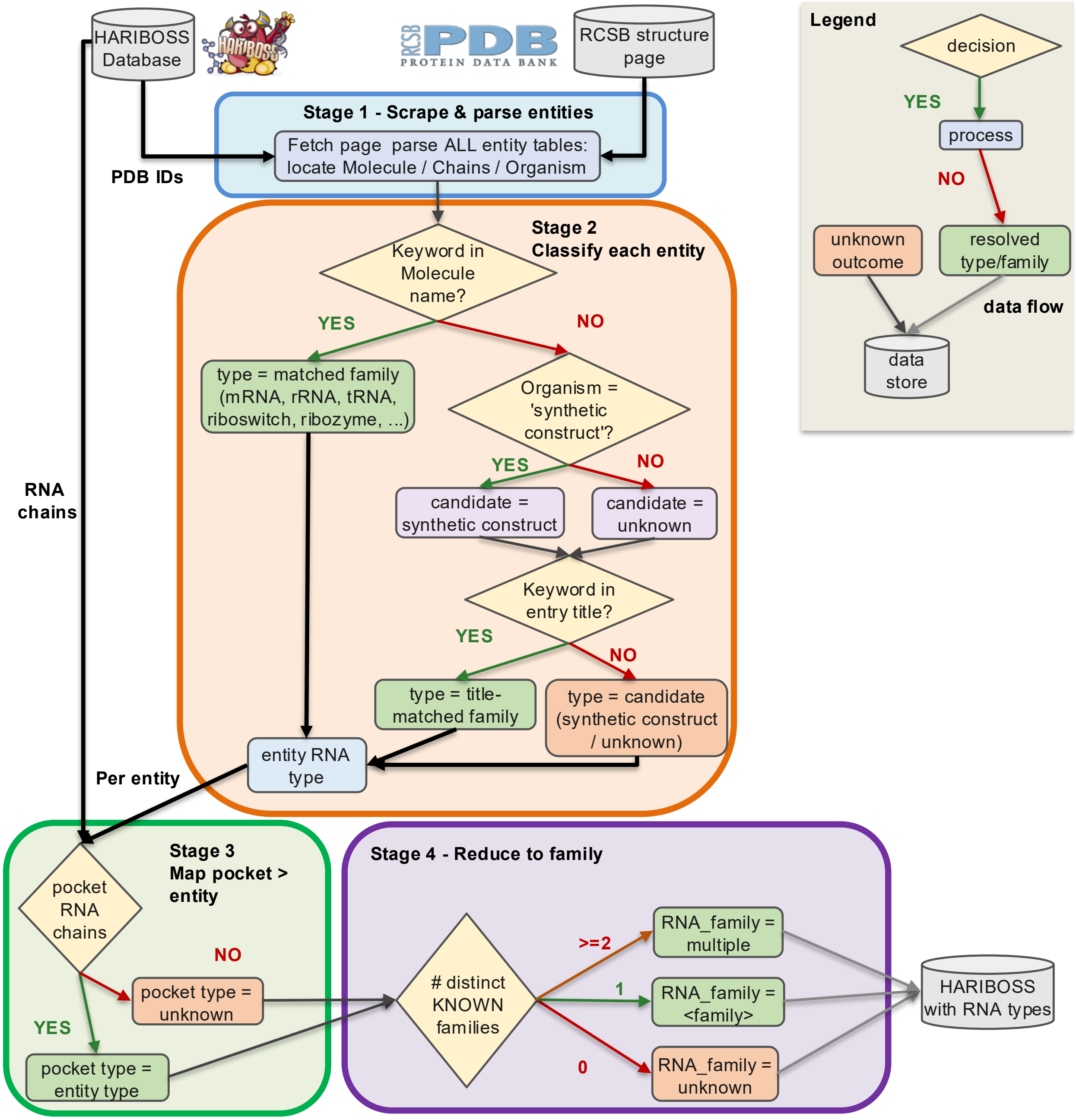


Figure S1: **RNA family annotation pipeline.** Schematic representation of the procedure used to annotate RNA chains family. Metadata are extracted from PDB entries (**Stage 1**, blue). Each RNA chain is associated to an RNA family using a rule-based keyword matching strategy (**Stage 2**, orange). RNA chains of residues belonging to binding pockets are assigned to their respective family (**Stage 3**, green). A global RNA family is assigned to the pocket (**Stage 4**, purple).

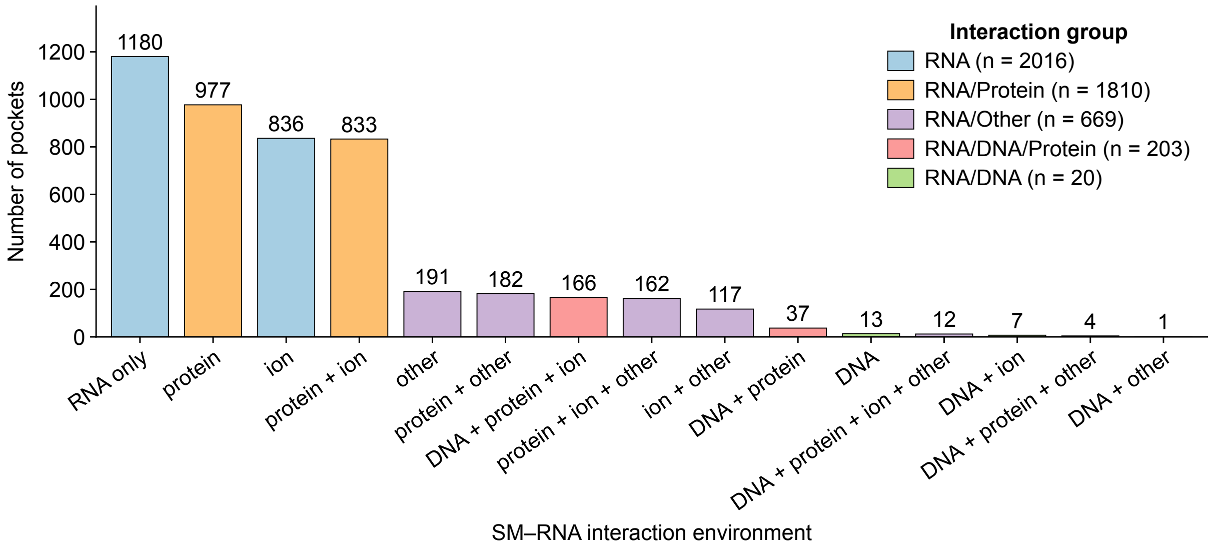


Figure S2: **Distribution of binding pockets across interaction environments in HARIBOSS ($n = 4718$).** Binding pockets are grouped according to their interaction environments: RNA (light blue, $n = 2016$), RNA/Protein (light orange, $n = 1810$), RNA/other (light purple, $n = 669$), RNA/DNA/Protein (light pink, $n = 203$), RNA/DNA (light green, $n = 20$).

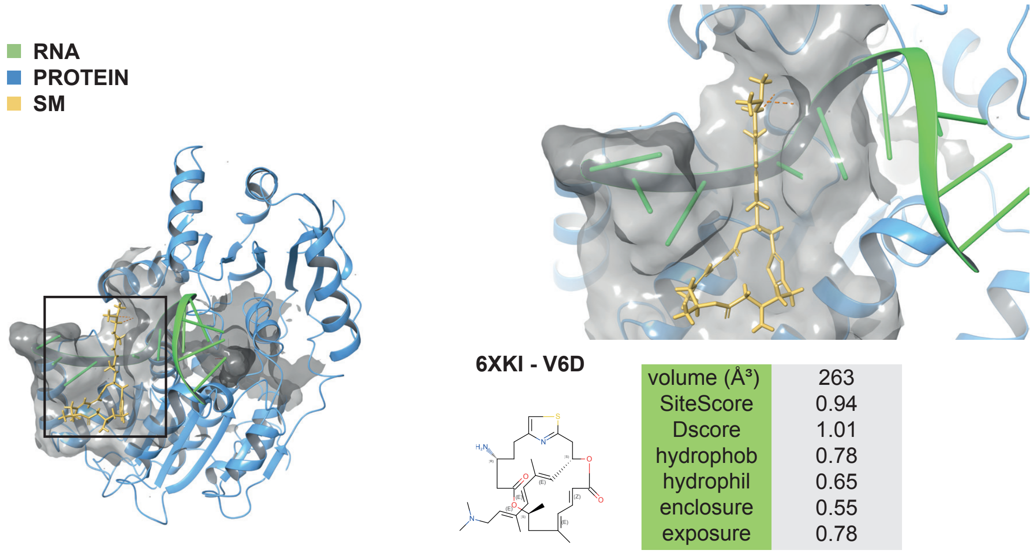


| | |
|---|---|
| volume (Å³) | 263 |
| SiteScore | 0.94 |
| Dscore | 1.01 |
| hydrophob | 0.78 |
| hydrophil | 0.65 |
| enclosure | 0.55 |
| exposure | 0.78 |

Figure S3: **Representative RNA/Protein binding pocket in PDB entry 6XKI.** Detailed view of the V6D ligand-binding pocket in PDB entry 6XKI[1]. The ligand binds at the interface between the eIF4A1 helicase protein (blue ribbon) and a short RNA polypurine sequence (green ribbon) as defined using the 6.0 Å cutoff to identify intermolecular contacts. This interaction stabilizes the closed conformation of eIF4A1, prevents RNA release and helicase turnover, inhibits ribosome scanning, and ultimately blocks translation initiation.

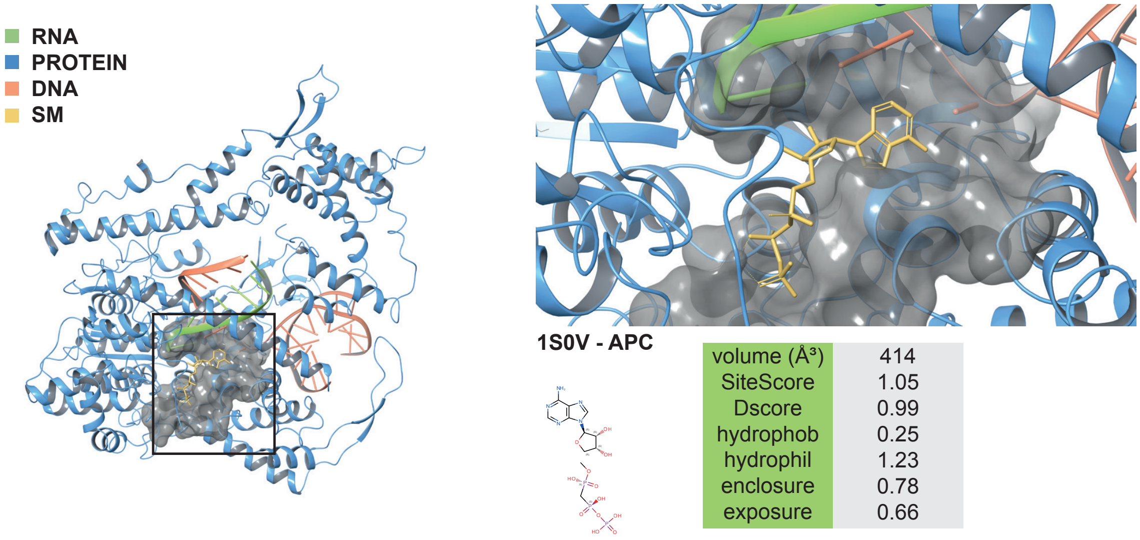


| | |
|---|---|
| volume (Å³) | 414 |
| SiteScore | 1.05 |
| Dscore | 0.99 |
| hydrophob | 0.25 |
| hydrophil | 1.23 |
| enclosure | 0.78 |
| exposure | 0.66 |

Figure S4: **Representative RNA/DNA/protein binding pocket in PDB entry 1S0V.** Detailed view of the APC ligand-binding pocket in PDB entry 1S0V[2]. The ligand simultaneously interacts with RNA (green ribbon), DNA (red ribbon), and protein (blue ribbon), as defined using the 6.0 Å cutoff to identify intermolecular contacts. This complex represents the elongation stage of transcription by T7 RNA polymerase, in which the DNA template strand base-pairs with the nascent RNA transcript to form a short RNA-DNA hybrid duplex stabilized by the polymerase.

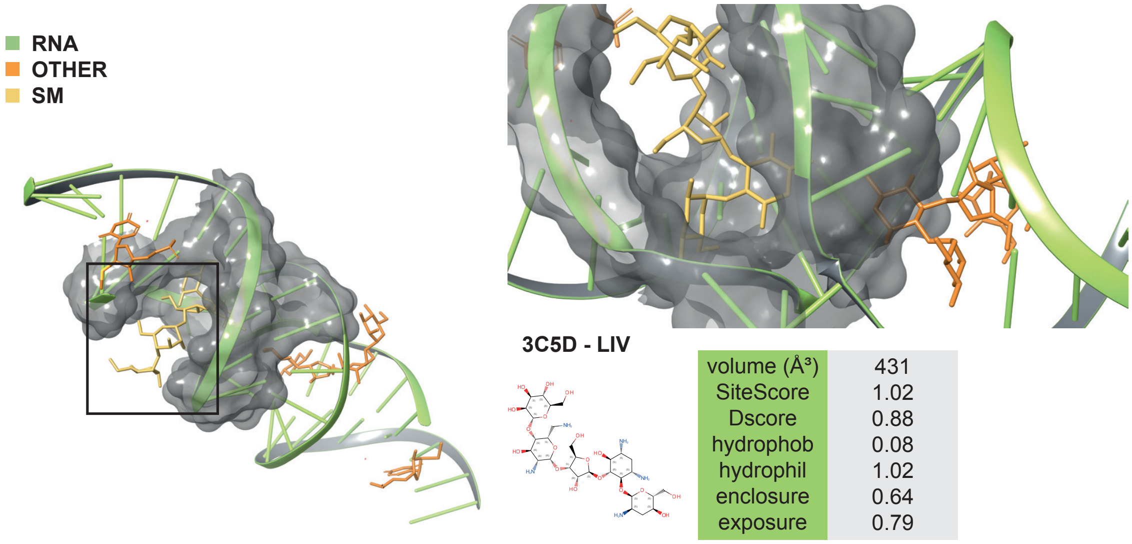


| | |
|---|---|
| volume (Å³) | 431 |
| SiteScore | 1.02 |
| Dscore | 0.88 |
| hydrophob | 0.08 |
| hydrophil | 1.02 |
| enclosure | 0.64 |
| exposure | 0.79 |

Figure S5: **Representative RNA/other binding pocket in PDB entry 3C5D.** Detailed view of the LIV ligand-binding pocket in PDB entry 3C5D[3]. The ligand (yellow) interacts with the RNA molecule (green ribbon) as well as with a second LIV molecule (orange), as defined using the 6.0 Å cutoff to identify intermolecular contacts. This structure represents the HIV-1 DIS extended duplex, in which two aminoglycoside molecules bind symmetrically to distorted RNA regions. Each ligand forms hydrogen-bonding and electrostatic interactions with the RNA phosphate backbone and exposed bases, while additional ligand-ligand contacts further stabilize the complex.

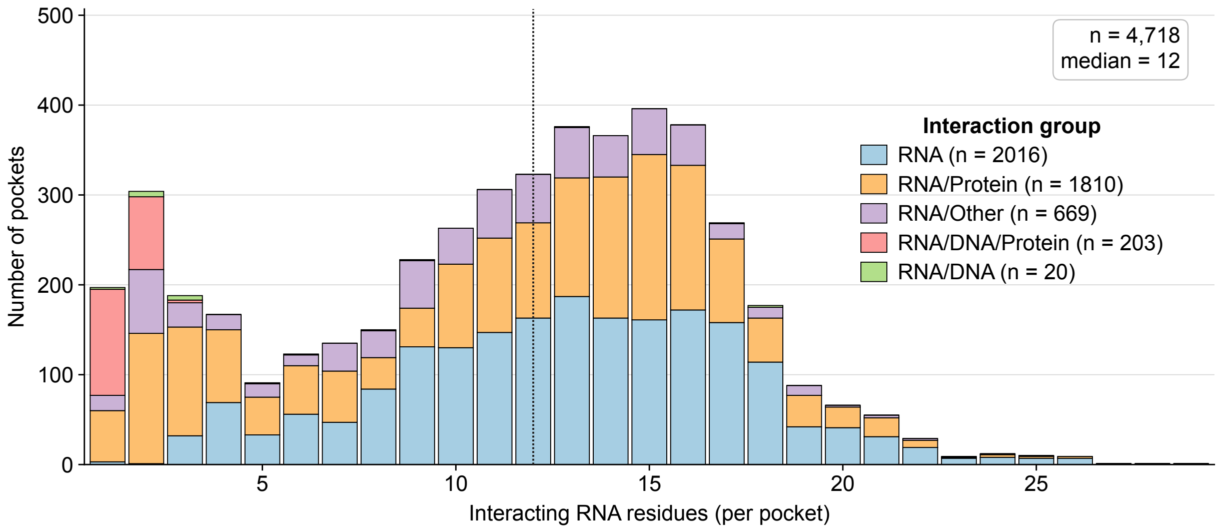


Figure S6: **Distribution of RNA nucleotides per binding pocket in the *redundant* HARIBOSS ($n = 4718$).** Bars are colored according to the interaction environment: RNA (light blue, $n = 2016$), RNA/Protein (light orange, $n = 1810$), RNA/Other (light purple, $n = 669$), RNA/DNA/Protein (light pink, $n = 203$), and RNA/DNA (light green, $n = 20$). The median number of RNA nucleotides per pocket is 12.

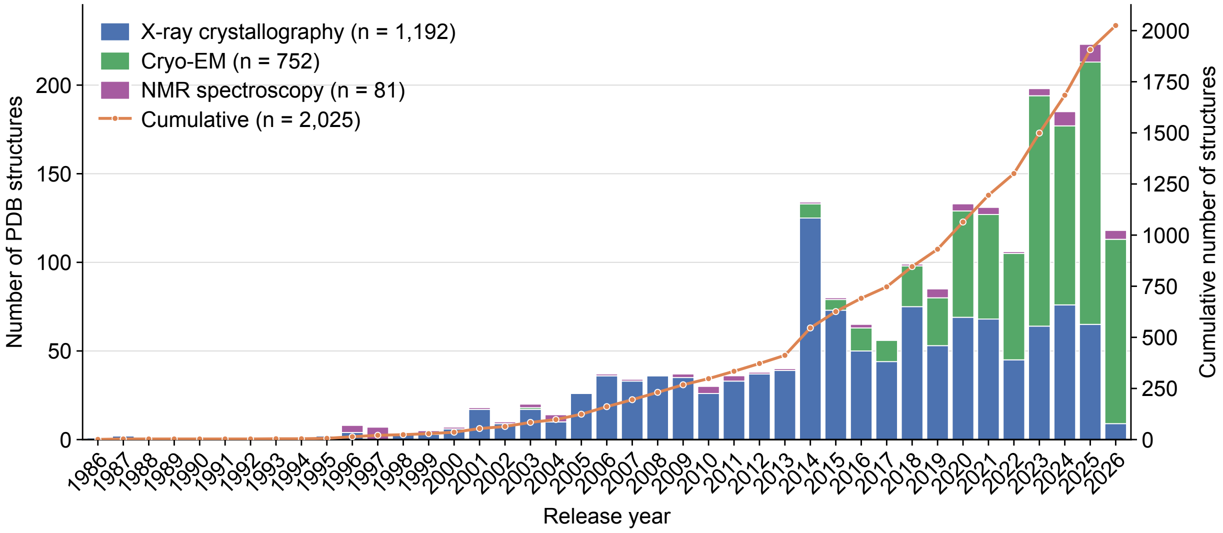


Figure S7: **Growth of the HARIBOSS database over time.** Numbers of PDB entries in the *redundant* HARIBOSS resolved by X-ray crystallography (dark blue, $n = 1192$), cryo-EM (green, $n = 752$) and NMR spectroscopy (dark purple, $n = 81$) as a function of released year, along with cumulative number of the structures (orange line).

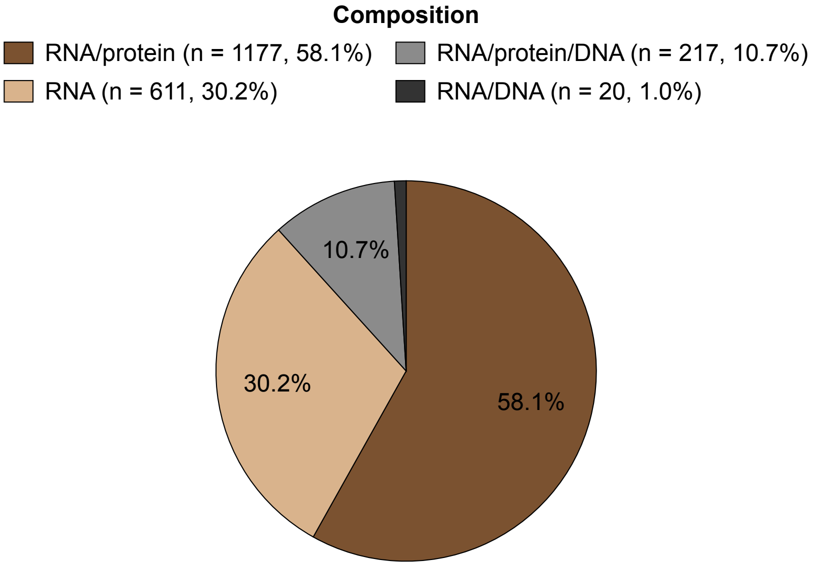


Figure S8: **Molecular composition of the *redundant* HARIBOSS database ($n = 2025$).** Complexes are grouped according to molecular composition: RNA (beige, $n = 611$), RNA/Protein (brown, $n = 1177$), RNA/Protein/DNA (grey, $n = 217$), and RNA/DNA (dark brown, $n = 20$).

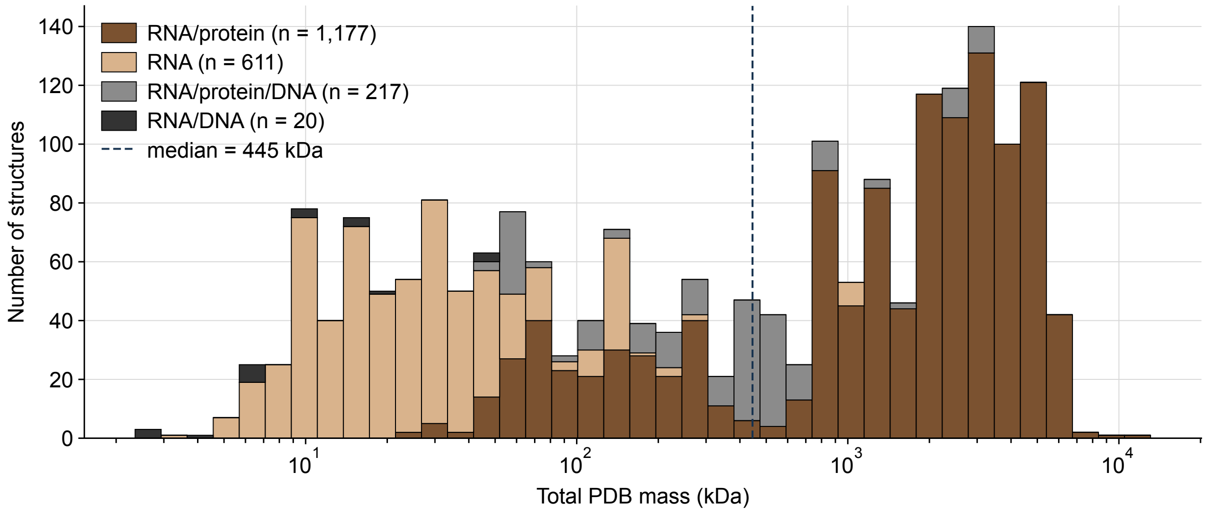


Figure S9: **Molecular mass distribution of the *redundant* HARIBOSS database ($n = 2025$).** Bars are colored according to the molecular composition of each complex: RNA (beige, $n = 611$), RNA/Protein (brown, $n = 1177$), RNA/Protein/DNA (grey, $n = 217$), and RNA/DNA (dark brown, $n = 20$).

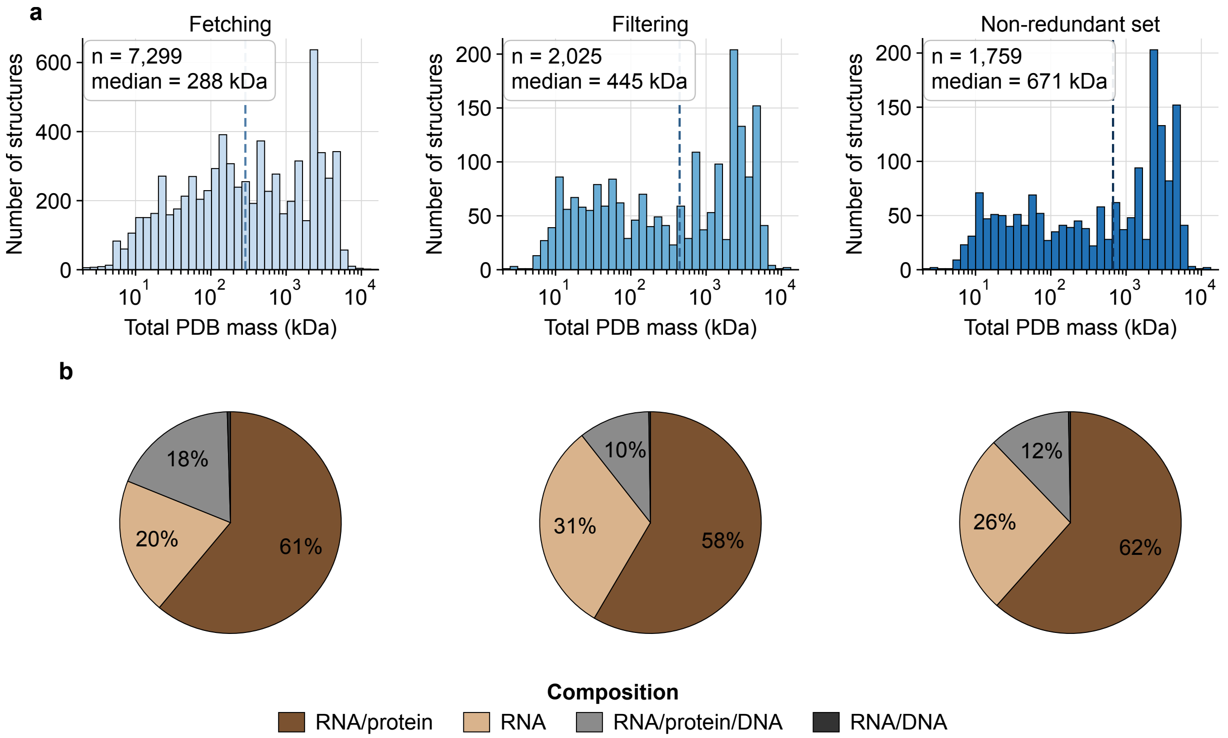


Figure S10: **Evolution of molecular mass distribution and molecular composition during the HARIBOSS construction steps. A.** Mass distribution of the RNA-small molecule complexes, fetching (light blue, $n = 7299$), filtering (blue, $n = 2025$), clustering (dark blue, $n = 1759$). **B.** Pie charts summarize the molecular composition of the dataset, with complexes colored according to their composition: RNA (beige), RNA/Protein (brown), RNA/Protein/DNA (grey), and RNA/DNA (dark brown).

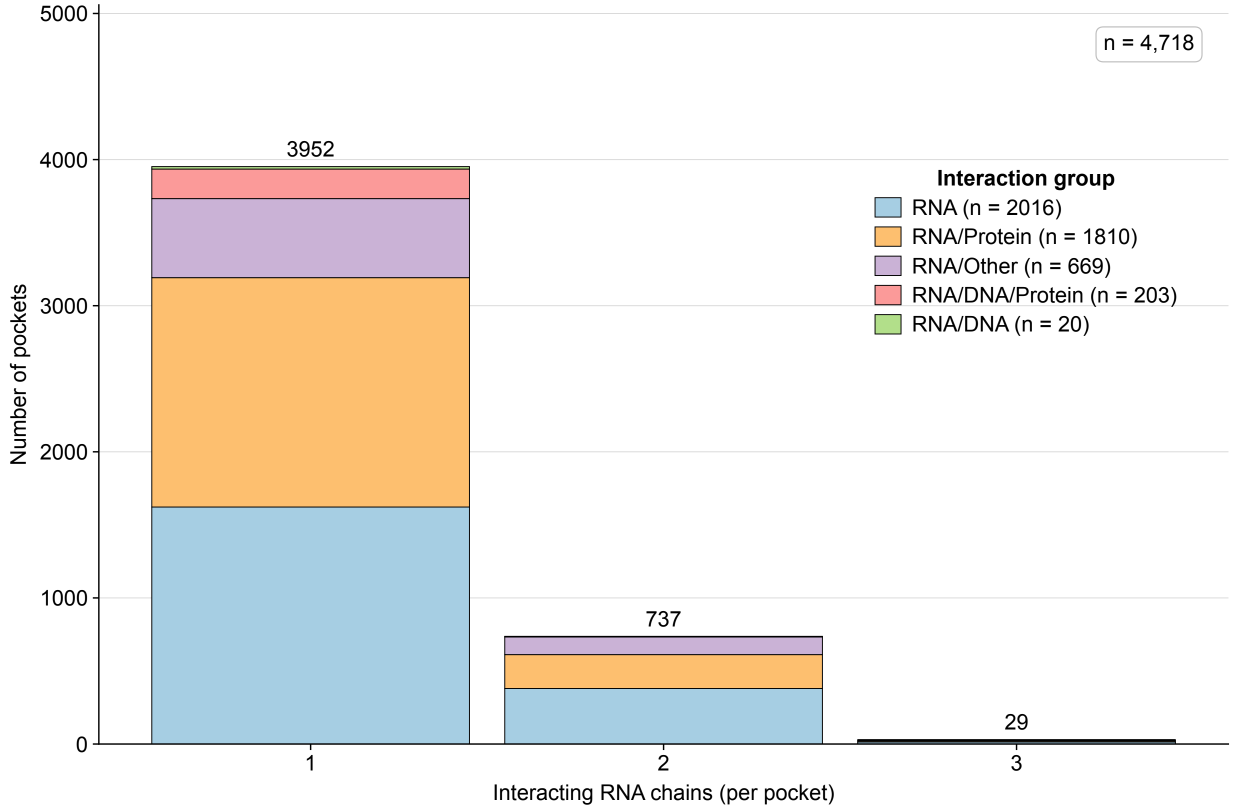


Figure S11: **Distribution of interacting RNA chains per binding pocket ($n = 4718$).** Distribution of binding pockets according to the number of interacting RNA chains the ligand interacts with. Bars are colored according to interaction environment: RNA (light blue), RNA/Protein (light orange), RNA/Other (light purple), RNA/DNA/Protein (light pink), and RNA/DNA (light green).

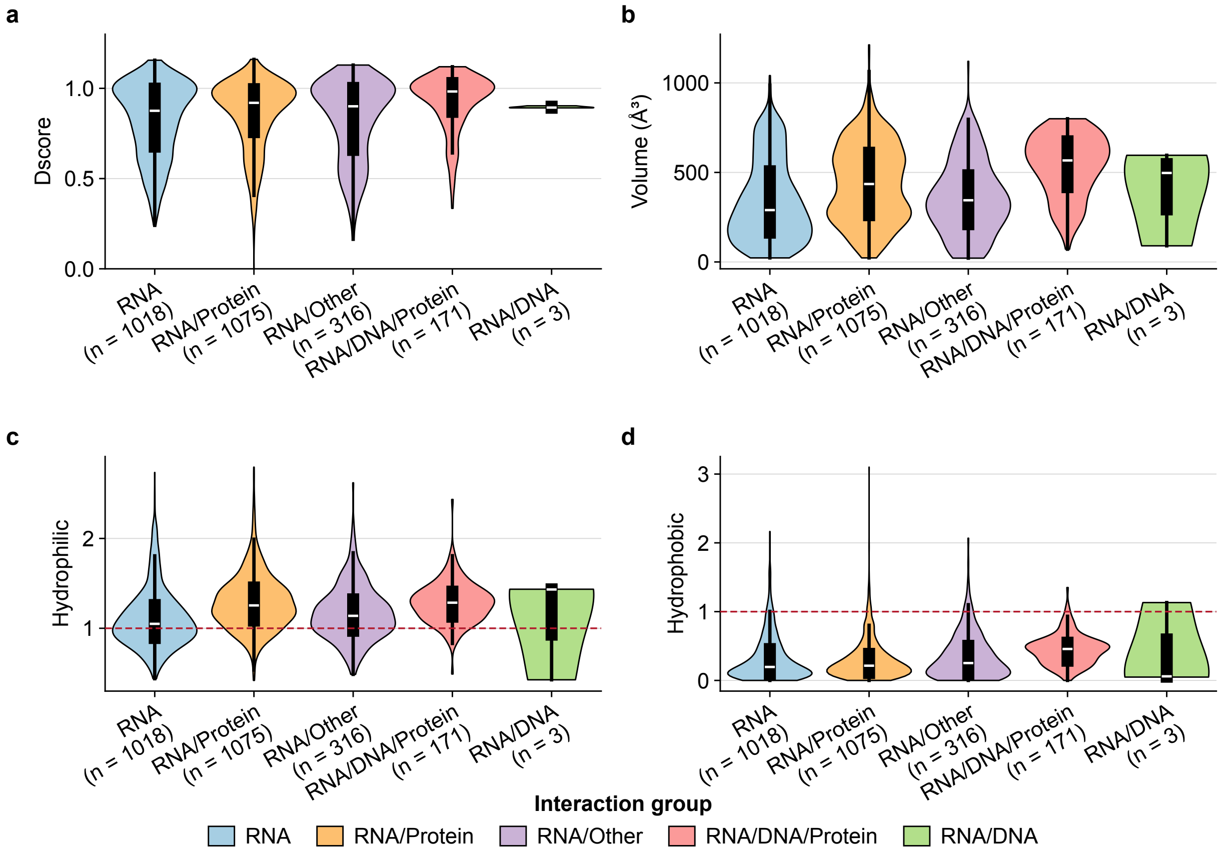


Figure S12: **SiteMap descriptors across interaction environments.** Comparison of SiteMap descriptors for curated, *non-redundant* HARIBOSS binding pockets ($n = 2583$) across the five interaction environments. **A & B.** Distribution of DScore and pocket volume. **C & D.** Distribution of pocket hydrophilicity and hydrophobicity.

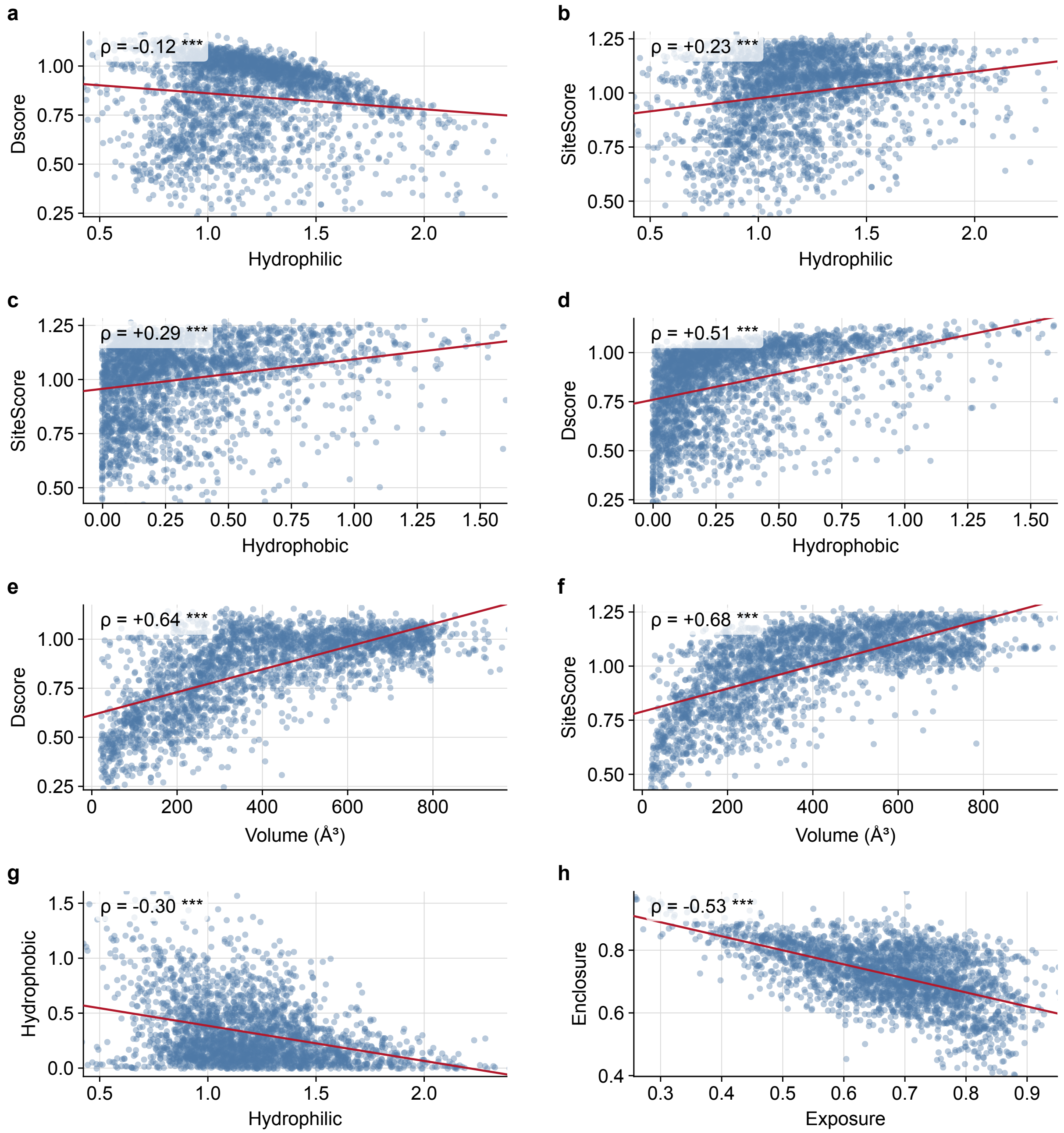


Figure S13: **Pairwise relationships between SiteMap descriptors.** Scatter plots showing pairwise relationships between SiteMap descriptors computed for curated, *non-redundant* HARIBOSS binding pockets ($n = 2583$). Spearman's rank correlation coefficient ($\rho$) is reported for each comparison. **A.** DScore versus hydrophilicity. **B.** SiteScore versus hydrophilicity. **C.** SiteScore versus hydrophobicity. **D.** DScore versus hydrophobicity. **E.** DScore versus pocket volume. **F.** SiteScore versus pocket volume. **G.** Hydrophobicity versus hydrophilicity. **H.** Enclosure versus exposure.

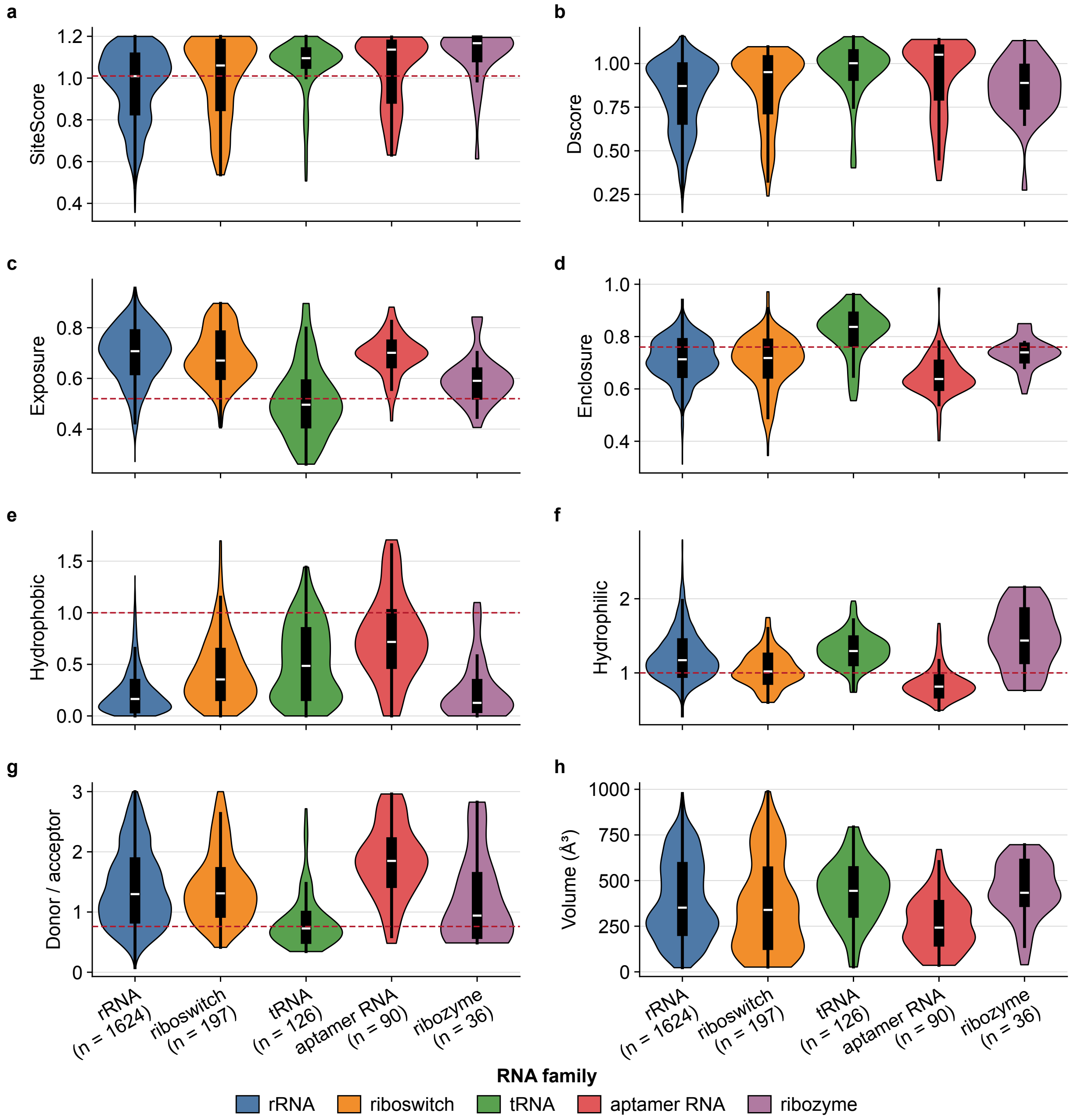


Figure S14: **SiteMap descriptors across major RNA families. A - H.** Distributions of SiteMap descriptors for curated, *non-redundant* HARIBOSS binding pockets grouped according to the five most represented RNA families: rRNA (blue, $n = 1624$), riboswitch (orange, $n = 197$), tRNA (green, $n = 126$), aptamer RNA (red, $n = 90$), and ribozyme (purple, $n = 36$).

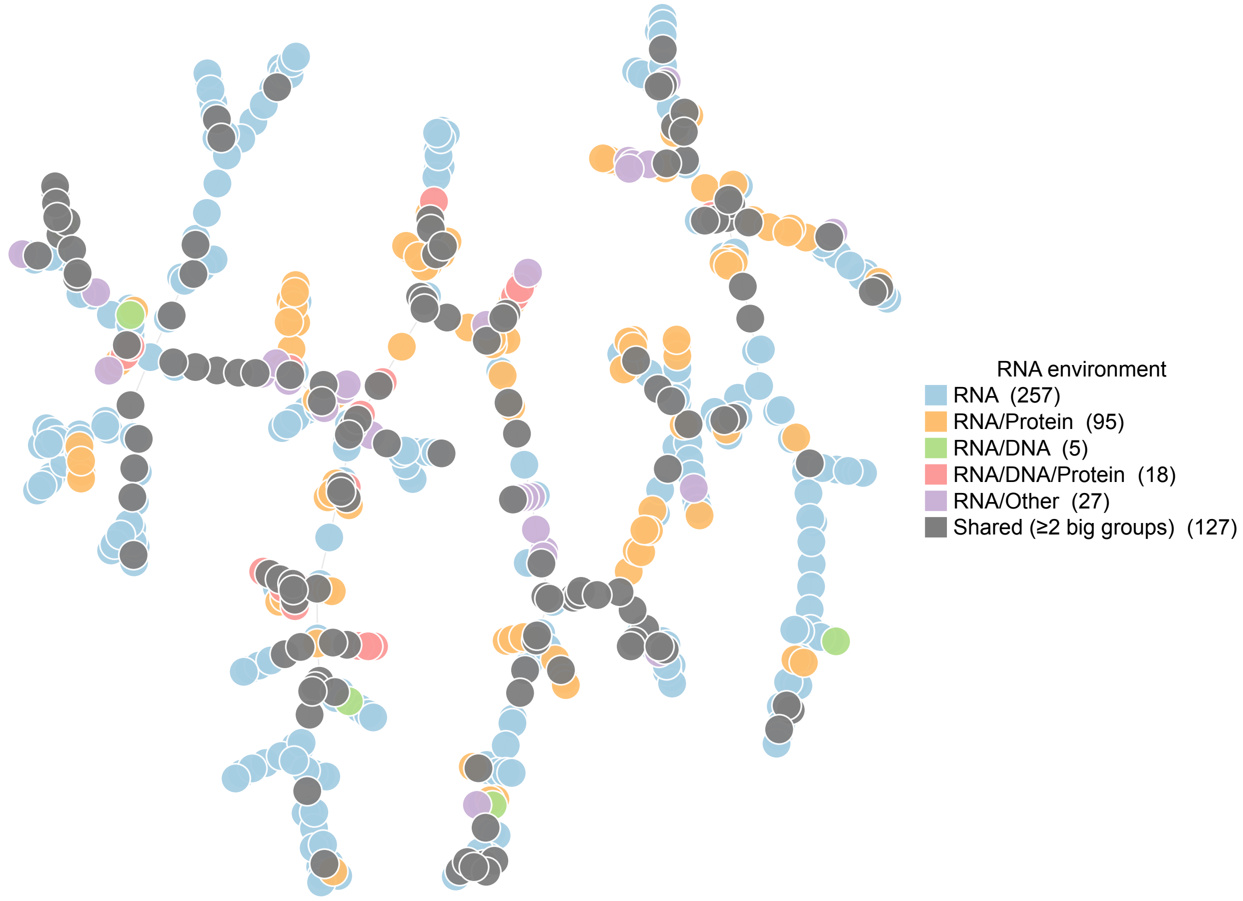


Figure S15: **TMAP visualization of ligand chemical space by interaction environment.** TMAP representation of the chemical space of HARIBOSS ligands ($n = 529$, 2 failed), colored according to their associated specific interaction environment: RNA (light blue, $n = 257$), RNA/Protein (light orange, $n = 95$), RNA/DNA (light green, $n = 5$), RNA/DNA/Protein (light red, $n = 18$), RNA/Other (light purple, $n = 27$), and ligands shared between multiple interaction environments (grey, $n = 127$).

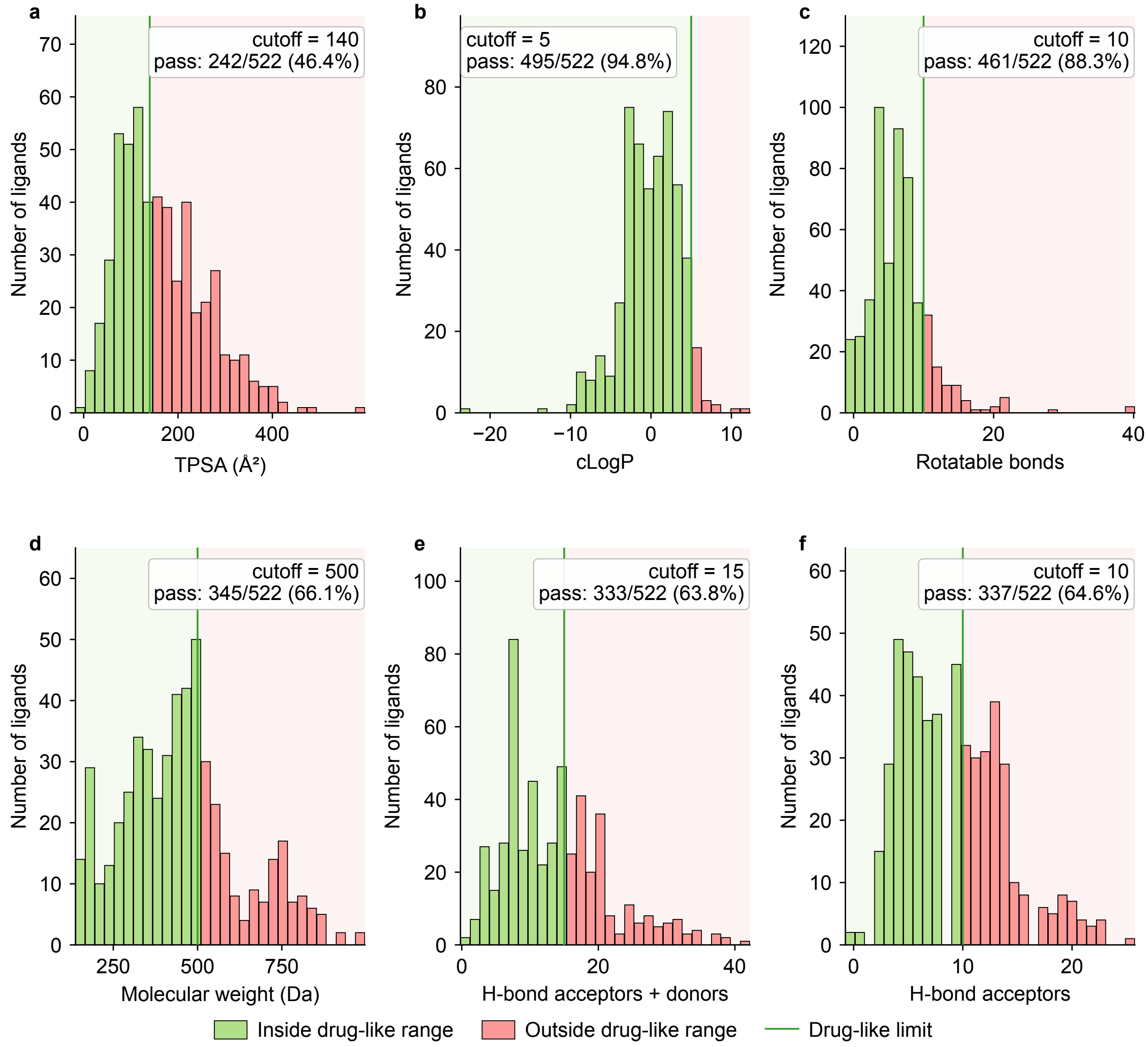


Figure S16: **Drug-like physicochemical properties of HARIBOSS ligands.** Distribution of key physicochemical properties for HARIBOSS ligands ($n = 522$, some failures exist in properties retrieval), evaluated against established drug-likeness criteria. **A.** Topological polar surface area (TPSA), reflecting molecular polarity and permeability. **B.** Octanol-water partition coefficient (LogP), indicating lipophilicity. **C.** Number of rotatable bonds, reflecting molecular flexibility. **D.** Molecular weight (Da). **E.** Total hydrogen-bonding capacity (HBA + HBD). **F.** Number of hydrogen-bond acceptors (HBA). Green shaded regions indicate commonly accepted Lipinski[4] and Veber[5] thresholds (MW $\leq$ 500 Da, LogP $\leq$ 5, HBD $\leq$ 5, HBA $\leq$ 10, TPSA $\leq$ 140 Å$^2$, and rotatable bonds $\leq$ 10), highlighting the fraction of ligands falling within classical drug-like chemical space.

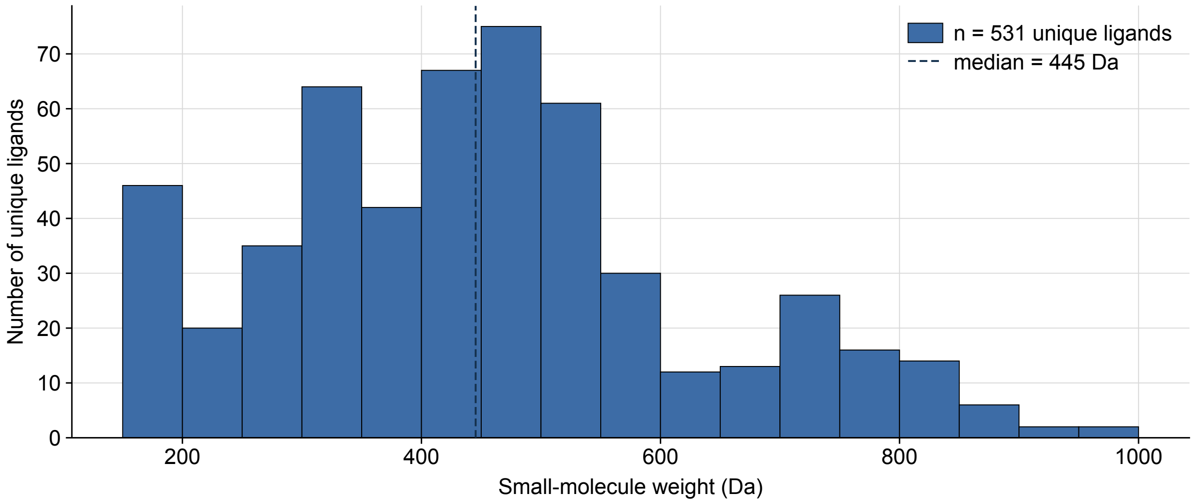


Figure S17: **Molecular weight distribution of HARIBOSS ligands.** Distribution of molecular weights for unique HARIBOSS ligands ($n = 531$). Molecular weights span a broad range, from 162.15 Da (ligand 29G) to 972.22 Da (ligand RU6), with a median value of 445 Da.

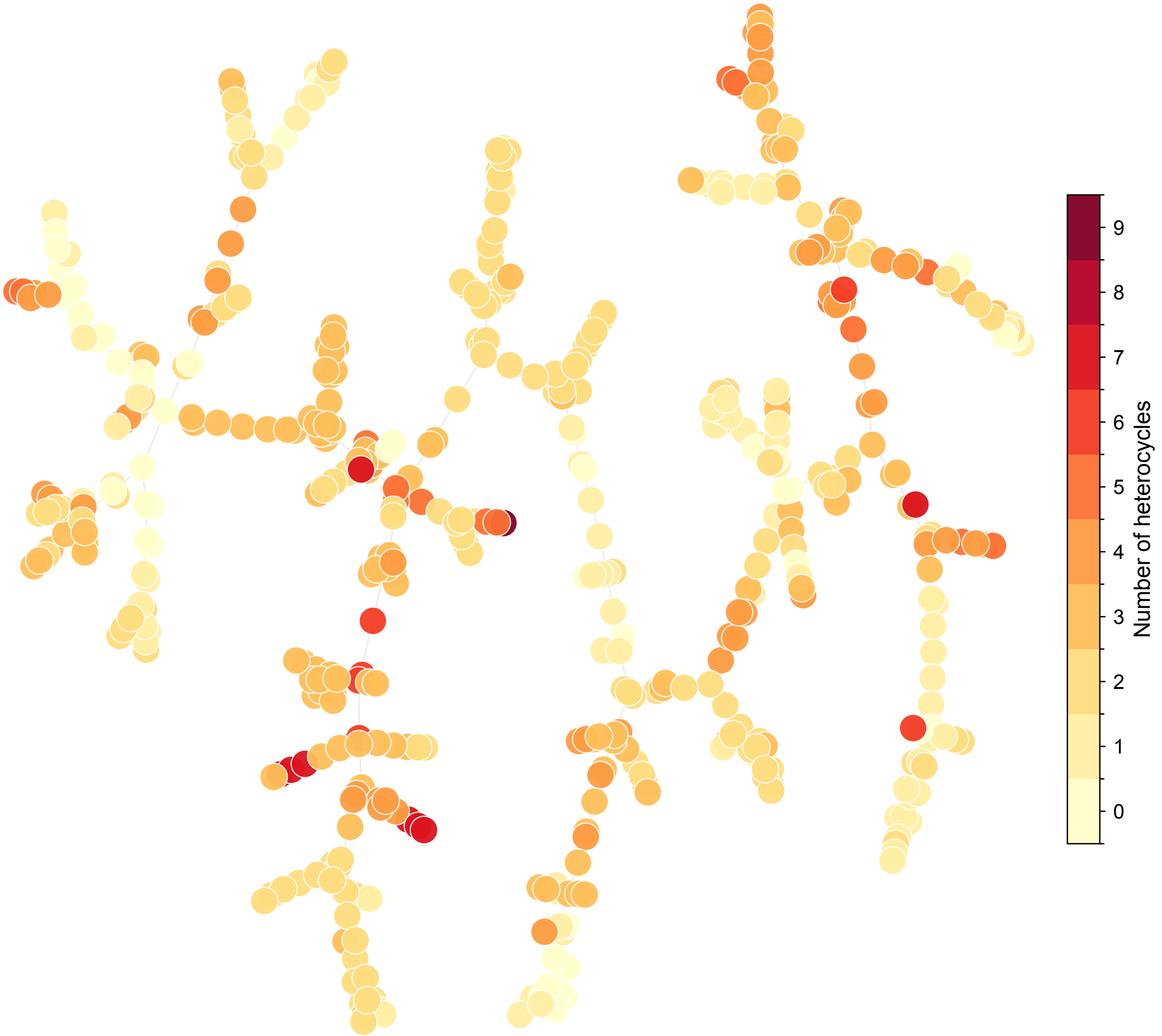


Figure S18: **TMAP visualization of ligand chemical space by heterocycle content.** TMAP representation of the chemical space of HARIBOSS ligands ($n$ = 529, 2 ligands failed), colored according to the number of heterocyclic rings present in each molecule. Colors range from light yellow (0 heterocycles) to dark red (9 heterocycles).

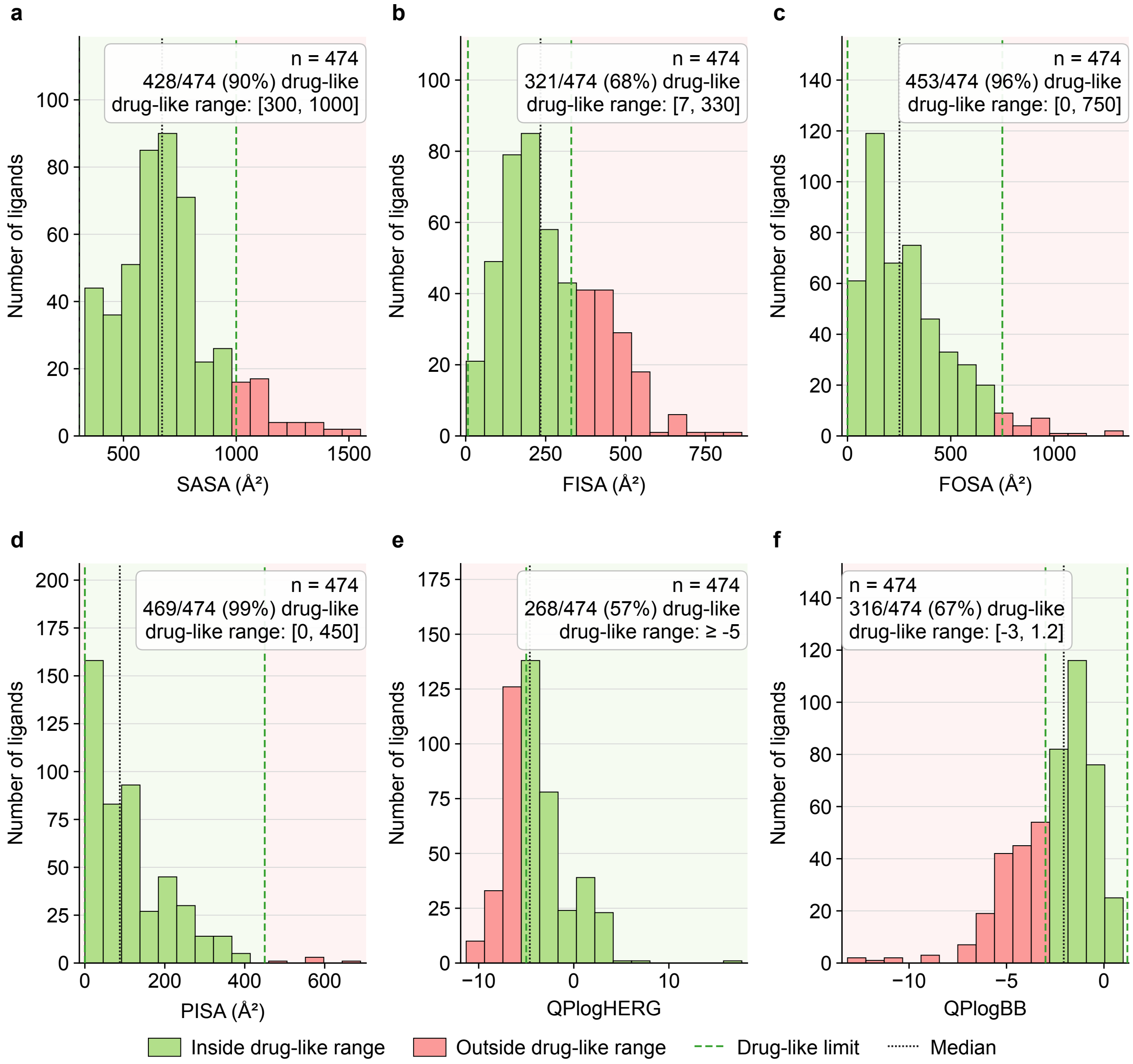


Figure S19: **QikProp molecular descriptors of HARIBOSS ligands.** Distribution of physicochemical and ADME-related properties computed with QikProp for HARIBOSS ligands ($n = 474$, QikProp failed for some ligands). **A.** Solvent-accessible surface area (SASA), reflecting overall molecular exposure to solvent. **B.** Hydrophilic solvent-accessible surface area (FISA). **C.** Hydrophobic solvent-accessible surface area (FOSA). **D.** $\pi$-component of the solvent-accessible surface area (PISA), associated with aromatic and conjugated systems. **E.** Predicted HERG $K^+$ channel inhibition (QPlogHERG), a proxy for potential cardiotoxicity. **F.** Predicted blood-brain barrier partition coefficient (QPlogBB), reflecting the ability of ligands to cross the blood-brain barrier.

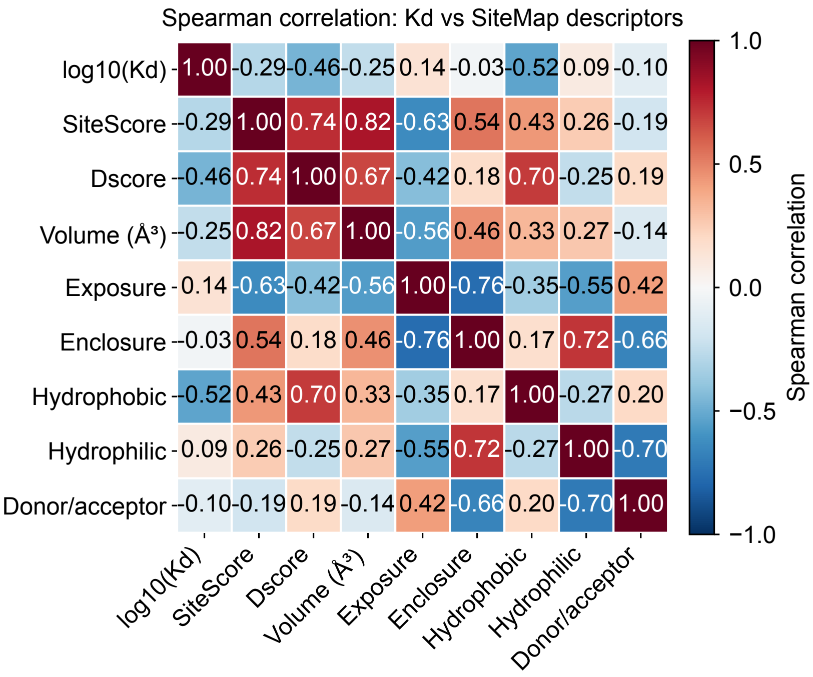


Figure S20: **Spearman correlation matrix of pocket descriptors.** Heatmap showing the pairwise Spearman rank correlation coefficients ($\rho$) between SiteMap descriptors computed for HARIBOSS binding pockets.

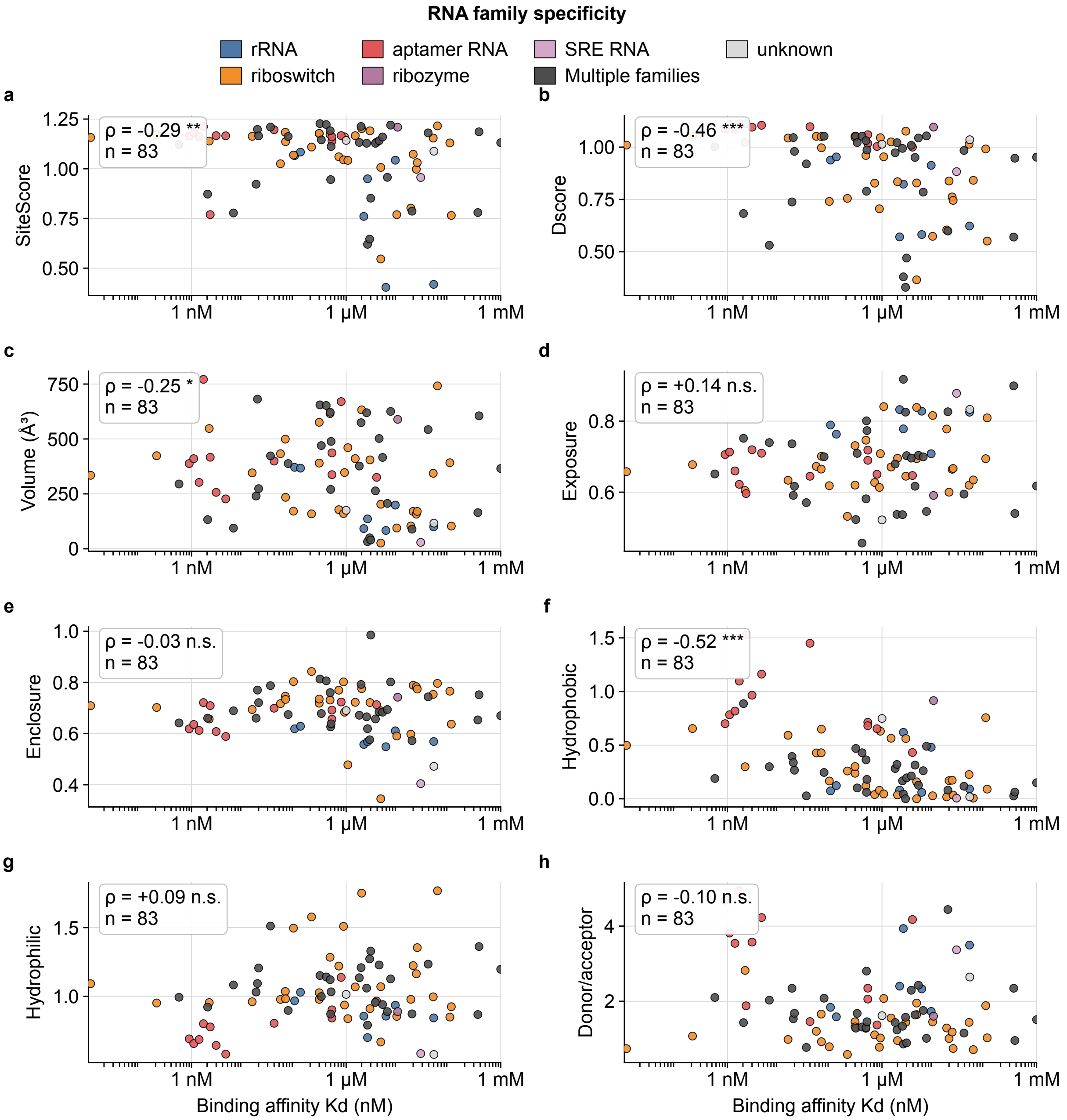


Figure S21: **Relationship between binding affinity and SiteMap descriptors.** Relationship between experimental binding affinity and SiteMap descriptors for RNA-small molecule complexes ($n$ = 83). Binding affinity is expressed as $K_d$ ($\log_{10}$ scale, nM) and grouped into three classes: very high affinity ($K_d$ < 1 nM), high affinity (1-$10^3$ nM), and moderate affinity ($10^3$-$10^6$ nM). **A.** SiteScore versus binding affinity. **B.** Pocket volume versus binding affinity. **C.** Pocket exposure versus binding affinity. **D.** Pocket enclosure versus binding affinity. **E.** Pocket hydrophobicity versus binding affinity. **F.** Hydrogen-bond donor/acceptor character (Don/Acc) versus binding affinity.

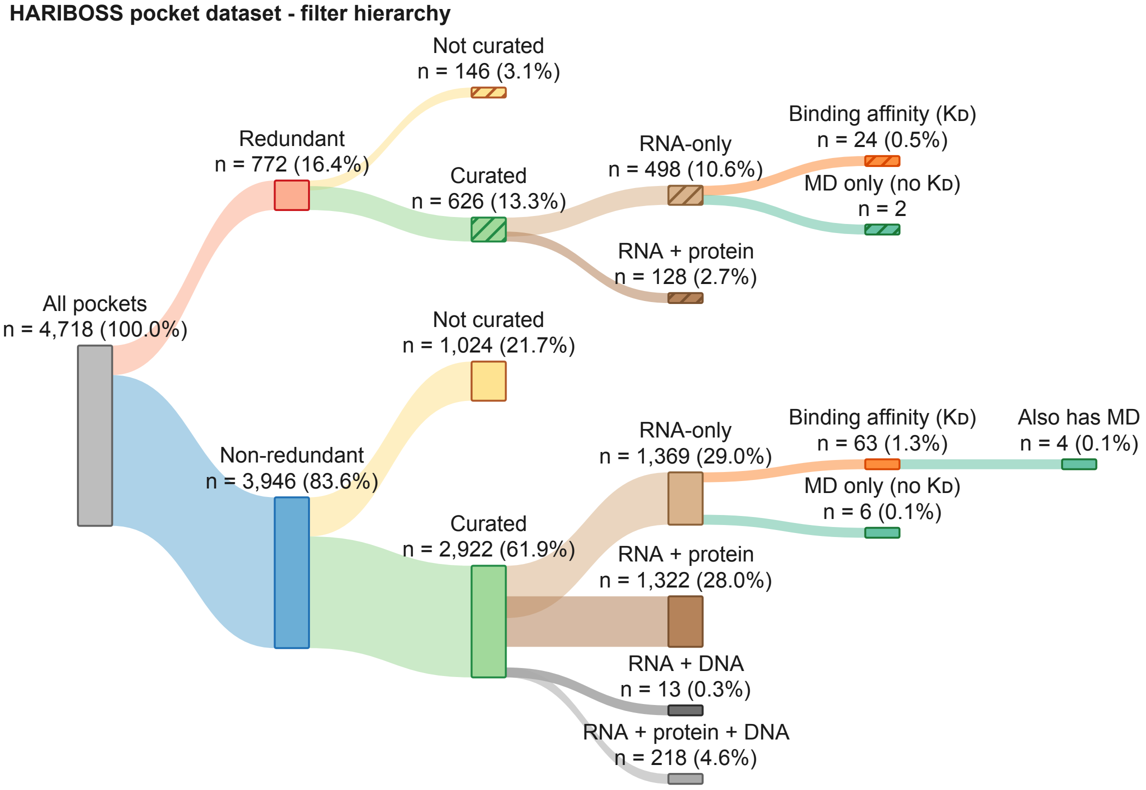


Figure S22: **Sankey diagram of the pockets in HARIBOSS.** Pockets are first divided between the *non-redundant* ones and *redundant* set. Within each set we split the curated set and we specify each interaction category. The subsets for which experimental binding-affinity data or molecular dynamics simulations are available are finally indicated.

Table S1: **Mapping of detailed interaction annotations to major interaction environments.** Each ligand-binding pocket is associated with one or more interacting molecular components in addition to RNA (protein, DNA, ions, or other molecules). These detailed interaction annotations are grouped into five major interaction environments used throughout HARIBOSS: RNA, RNA/Protein, RNA/DNA, RNA/DNA/Protein, and RNA/Other.

| **Interacting component in addition to RNA** | **Assigned group** |
|---|---|
| `-` | RNA |
| `ion` | RNA |
| `protein` | RNA/Protein |
| `protein`, `ion` | RNA/Protein |
| `DNA` | RNA/DNA |
| `DNA`, `ion` | RNA/DNA |
| `DNA`, `protein` | RNA/DNA/protein |
| `DNA`, `protein`, `ion` | RNA/DNA/protein |
| `other` | RNA/Other |
| `ion`, `other` | RNA/Other |
| `protein`, `other` | RNA/Other |
| `protein`, `ion`, `other` | RNA/Other |
| `DNA`, `other` | RNA/Other |
| `DNA`, `protein`, `other` | RNA/Other |
| `DNA`, `protein`, `ion`, `other` | RNA/Other |

Table S2: **Overview of the major RNA families.** Summary of the principal RNA families identified in HARIBOSS, including their primary biological role and key functional characteristics[6].

| RNA family | Primary role | Key features / function |
|---|---|---|
| mRNA | Coding | Template for protein synthesis<br>Contains codons translated by ribosomes |
| rRNA | Structural / catalytic | Core component of ribosome |
| tRNA | Adapter | Transfers amino acids<br>Anticodon pairs with mRNA codons |
| Riboswitch | Regulatory | Ligand-binding RNA<br>Regulates gene expression via conformational change |
| SRE RNA | Regulatory | Splicing regulatory elements |
| snRNA | Regulatory / catalytic | Spliceosome components<br>Mediate pre-mRNA splicing |
| Ribozyme | Catalytic | RNA enzyme<br>Performs cleavage or ligation reactions |
| Telomerase RNA | Template | Provides template for telomere extension |
| TAR RNA | Viral regulatory | HIV RNA element<br>Binds Tat protein to regulate transcription |
| Aptamer RNA | Ligand-binding | Binds specific ligands with high affinity<br>Natural or SELEX-derived structures |
| Synthetic construct | Engineered | Artificial RNA sequences<br>Designed for structural or functional studies |
| RNA/DNA hybrid | Hybrid structure | RNA–DNA duplex |

Table S3: **Common drug-likeness rules used in medicinal chemistry.** Summary of widely used physicochemical criteria for assessing drug-like molecules, including Lipinski's Rule of Five[4], Veber's rules[5], and Pfizer's 3/75 rule[7].

| Rule Set | Property | Condition | Primary Goal |
|---|---|---|---|
| **Lipinski's Rule of 5[4]** | Molecular Weight (MW) | $\leq 500$ Da | Oral Absorption |
| | LogP | $\leq 5$ | |
| | H-Bond Donors (HBD) | $\leq 5$ | |
| | H-Bond Acceptors (HBA) | $\leq 10$ | |
| **Veber's Rules[5]** | Rotatable Bonds | $\leq 10$ | Bioavailability |
| | Topological Polar Surface Area (TPSA) | $\leq 140\,\mathring{A}^2$ | |
| **Pfizer's 3/75 Rule[7]** | LogP | $> 3$ | Toxicity Safety |
| | Topological Polar Surface Area (TPSA) | $< 75\,\mathring{A}^2$ | |

Table S4: **List of abbreviations.** These abbreviations are used throughout the manuscript and are important to understand tools and metrics that have been used.

| **Abbreviation** | **Definition** |
|---|---|
| PDB | Protein Data Bank |
| RNP | Ribonucleoprotein |
| NR | Non-redundant |
| IFE | Integrated Functional Element |
| MDDB | Molecular Dynamics Data Bank |
| BGSU | Bowling Green State University |
| TPSA | Topological Polar Surface Area |
| HBA | Hydrogen-Bond Acceptor |
| HBD | Hydrogen-Bond Donor |
| SASA | Solvent-Accessible Surface Area |
| FISA | Hydrophilic Component of the Solvent-Accessible Surface Area |
| FOSA | Hydrophobic Component of the Solvent-Accessible Surface Area |
| PISA | $\pi$ Component of the Solvent-Accessible Surface Area |
| WPSA | Weakly Polar Component of the Solvent-Accessible Surface Area |

Table S5: **QikProp molecular descriptors and recommended ranges.** Description of the physicochemical and ADME-related descriptors computed with QikProp, together with the corresponding recommended value ranges.

| **Property** | **Description** | **Range** |
|---|---|---|
| SASA | Total solvent accessible surface area in $Å^2$ | 300.0 - 1000.0 |
| FOSA | Hydrophobic component of the SASA | 0.0 - 750.0 |
| FISA | Hydrophilic component of the SASA | 7.0 - 330.0 |
| PISA | $\pi$ component of the SASA (carbon & attached hydrogen) | 0.0 - 450.0 |
| WPSA | Weakly polar component of the SASA (halogens, P, & S) | 0.0 - 175.0 |
| Volume | Total solvent-accessible volume in $Å^3$ using a 1.4 Å probe | 500.0 - 2000.0 |
| QPlogHERG | Predicted $IC_{50}$ for blockage of HERG $K^+$ channels | Concern below $-5$ |
| QPPCaco | Predicted apparent Caco-2 cell permeability (nm/s) | $< 25$ poor $> 500$ great |
| QPlogBB | Predicted brain/blood partition coefficient | $-3.0$ – 1.2 |

Table S6: **Biological cofactors in HARIBOSS.** List of biological cofactors (abbreviation and full name) present in HARIBOSS and kept during binding pocket analysis.

| **Abbreviation** | **Full name** |
|---|---|
| ATP | Adenosine triphosphate |
| GTP | Guanosine triphosphate |
| NAD | Nicotinamide adenine dinucleotide |
| FAD | Flavin adenine dinucleotide |
| CoA | Coenzyme A |
| SAM | S-Adenosyl-L-methionine |
| THF | Tetrahydrofolate |
| – | Heme |
| PLP | Pyridoxal 5′-phosphate |
| TPP | Thiamine pyrophosphate |
| – | Biotin |
| FMN | Flavin mononucleotide |

Table S7: **Curated list of ligands to remove for druggability analyses.** List of the ligands (abbreviation and full name) in HARIBOSS not commonly considered as drugs. Their associated binding pockets were not considered when analyzing druggability properties.

| **Abbreviation** | **Full name** |
|---|---|
| SPM | Spermine |
| SPK | Fully Protonated form of the Spermine |
| EPE | 4-(2-Hydroxyethyl)-1-piperazineethanesulfonic acid (HEPES) |
| MES | 2-(N-Morpholino)ethanesulfonic acid |
| CIT | Citrate |
| BTB | Bis-tris buffer |
| TAM | Tris(hydroxymethyl)aminomethane (Tris) |
| PG4 | Tetraethylene glycol |
| 1PE | Pentaethylene glycol |
| PGV | Phosphatidylglycerol |
| PEV | Phosphatidylethanolamine |
| GLC | D-Glucose |
| CYY | Cysteine |
| 2TB | 1,3-diamino-4,5,6-Trihydroxy-cyclohexane |

Table S8: **Composition statistics of the HARIBOSS database, curated pockets excluded.** Numbers of RNA-SM complexes, pockets and ligands when removing data associated to our curated list of ligands. Data are categorized by structural composition of the binding pocket (RNA-only versus multi-biomolecule interfaces) and dataset redundancy.

| Structural Category | Dataset Type | RNA-SM complexes | Pockets | Ligands |
|---|---|---|---|---|
| **RNA-only** | Redundant | 975 | 1598 | 352 |
| | Non-redundant | 777 | 1201 | 344 |
| **RNA (with or w/o partner)** | Redundant | 1862 | 3548 | 518 |
| | Non-redundant | 1621 | 2922 | 504 |

Table S9: **SiteScore and Dscore statistics among different interaction environments.** Proportion of curated, *non-redundant* binding pockets satisfying commonly used SiteMap thresholds (SiteScore $\geq$ 0.8, DScore $\geq$ 0.98, and 0.83 $\leq$ DScore $<$ 0.98) for each interaction environment. Percentages are calculated relative to the number of SiteMap evaluated pockets within each group.

| **Interaction Group** | **SiteScore $\geq$ 0.8** | **DScore $\geq$ 0.98** | **0.83 $\leq$ DScore $<$ 0.98** |
|---|---|---|---|
| RNA | 855 (84.0%) | 311 (30.6%) | 262 (25.7%) |
| RNA/Protein | 907 (84.4%) | 328 (30.5%) | 378 (35.2%) |
| RNA/Other | 246 (77.8%) | 105 (33.2%) | 86 (27.2%) |
| RNA/DNA/Protein | 154 (90.1%) | 89 (52.0%) | 46 (26.9%) |
| RNA/DNA | 3 (100%) | 0 (0%) | 3 (100%) |

Table S10: **SiteMap descriptors statistics across interaction environments.** Median, mean, minimum, and maximum values of SiteMap descriptors computed for curated, *non-redundant* HARIBOSS binding pockets grouped according to their interaction environment.

| Feature / RNA environment | Median | Mean | Max | Min |
|---|---|---|---|---|
| **SiteScore** | | | | |
| RNA ($n = 1018$) | 1.064 | 1.0141 | 1.269 | 0.403 |
| RNA/Protein ($n = 1075$) | 1.054 | 1.0012 | 1.275 | 0.357 |
| RNA/Other ($n = 316$) | 0.996 | 0.9494 | 1.240 | 0.398 |
| RNA/DNA/Protein ($n = 171$) | 1.066 | 1.0130 | 1.228 | 0.609 |
| RNA/DNA ($n = 3$) | 1.150 | 1.0982 | 1.164 | 0.981 |
| **Dscore** | | | | |
| RNA | 0.876 | 0.8254 | 1.156 | 0.241 |
| RNA/Protein | 0.920 | 0.8544 | 1.160 | -0.061 |
| RNA/Other | 0.900 | 0.8294 | 1.129 | 0.164 |
| RNA/DNA/Protein | 0.983 | 0.9244 | 1.120 | 0.335 |
| RNA/DNA | 0.893 | 0.8961 | 0.903 | 0.892 |
| **Exposure** | | | | |
| RNA | 0.712 | 0.7124 | 0.956 | 0.408 |
| RNA/Protein | 0.667 | 0.6430 | 0.955 | 0.233 |
| RNA/Other | 0.690 | 0.6773 | 0.908 | 0.185 |
| RNA/DNA/Protein | 0.604 | 0.6128 | 0.901 | 0.406 |
| RNA/DNA | 0.617 | 0.6583 | 0.774 | 0.584 |
| **Enclosure** | | | | |
| RNA | 0.692 | 0.6817 | 0.985 | 0.312 |
| RNA/Protein | 0.760 | 0.7514 | 0.963 | 0.313 |
| RNA/Other | 0.713 | 0.7100 | 0.997 | 0.327 |
| RNA/DNA/Protein | 0.812 | 0.8070 | 0.954 | 0.644 |
| RNA/DNA | 0.697 | 0.6512 | 0.718 | 0.538 |
| **Don/Acc** | | | | |
| RNA | 1.740 | 1.9842 | 4.997 | 0.303 |
| RNA/Protein | 1.047 | 1.2525 | 4.990 | 0.075 |
| RNA/Other | 1.191 | 1.4284 | 4.960 | 0.057 |
| RNA/DNA/Protein | 0.763 | 0.8407 | 2.525 | 0.281 |
| RNA/DNA | 1.715 | 2.3902 | 3.915 | 1.541 |

Table S11: **Representative PDB entries for major RNA families.** Examples of PDB entries containing at least one RNA chain representative of each RNA family identified in HARIBOSS.

| **RNA Type** | **PDB id** |
|---|---|
| riboswitch | 2GDI |
| TAR RNA | 1LVJ |
| SRE RNA | 1EI2 |
| ribozyme | 7YCG |
| tRNA | 7K98 |
| rRNA | 2HHH |
| telomerase RNA | 3MIJ |
| snRNA | 8CF2 |
| RNA/DNA hybrid | 7M7M |
| aptamer RNA | 6UP0 |
| mRNA | 6T0V |
| synthetic construct | 7OM7 |
| unknown | 7OJN |

Table S12: **Distribution of RNA families across ligand-binding pockets.** Distribution ofbinding pockets ($n = 4718$) among RNA families in *redundant* HARIBOSS. RNA family annotations were successfully assigned to 4406 of 4718 pockets (93.4%).

| **RNA family** | **Pockets** | **%** |
|---|---|---|
| rRNA | 2873 | 60.9 |
| riboswitch | 444 | 9.4 |
| synthetic construct | 336 | 7.1 |
| unknown | 312 | 6.6 |
| aptamer | 175 | 3.7 |
| tRNA | 156 | 3.3 |
| ribozyme | 156 | 3.3 |
| multiple | 107 | 2.3 |
| RNA/DNA hybrid | 61 | 1.3 |
| TAR RNA | 41 | 0.9 |
| mRNA | 31 | 0.7 |
| sre RNA | 11 | 0.2 |
| telomerase RNA | 10 | 0.2 |
| snRNA | 5 | 0.1 |
| **Total** | 4718 | 100 |

Table S13: **SiteMap descriptor statistics across major RNA families.** Median, mean, minimum, and maximum values of selected SiteMap descriptors (SiteScore, DScore, Exposure, Enclosure, and Don/Acc) computed for curated, *non-redundant* binding pockets belonging to rRNA, riboswitch, tRNA, aptamer RNA and ribozyme.

| **Feature** | **Median** | **Mean** | **Max** | **Min** |
|---|---|---|---|---|
| **SiteScore** | | | | |
| rRNA ($n = 1624$) | 1.026 | 0.988 | 1.269 | 0.357 |
| riboswitch ($n = 197$) | 1.122 | 1.038 | 1.238 | 0.537 |
| tRNA ($n = 126$) | 1.096 | 1.066 | 1.222 | 0.507 |
| aptamer RNA ($n = 90$) | 1.142 | 1.055 | 1.243 | 0.631 |
| ribozyme ($n = 36$) | 1.177 | 1.129 | 1.228 | 0.613 |
| **Dscore** | | | | |
| rRNA | 0.871 | 0.817 | 1.156 | -0.062 |
| riboswitch | 0.951 | 0.867 | 1.096 | 0.241 |
| tRNA | 1.002 | 0.957 | 1.154 | 0.402 |
| aptamer RNA | 1.051 | 0.935 | 1.138 | 0.330 |
| ribozyme | 0.889 | 0.859 | 1.132 | 0.275 |
| **Exposure** | | | | |
| rRNA | 0.707 | 0.698 | 0.956 | 0.272 |
| riboswitch | 0.671 | 0.686 | 0.896 | 0.408 |
| tRNA | 0.496 | 0.511 | 0.896 | 0.262 |
| aptamer RNA | 0.701 | 0.693 | 0.881 | 0.432 |
| ribozyme | 0.591 | 0.595 | 0.842 | 0.406 |
| **Enclosure** | | | | |
| rRNA | 0.713 | 0.714 | 0.939 | 0.312 |
| riboswitch | 0.718 | 0.704 | 0.971 | 0.345 |
| tRNA | 0.837 | 0.816 | 0.961 | 0.555 |
| aptamer RNA | 0.638 | 0.650 | 0.985 | 0.402 |
| ribozyme | 0.740 | 0.736 | 0.849 | 0.581 |
| **Don/Acc** | | | | |
| rRNA | 1.402 | 1.628 | 4.990 | 0.075 |
| riboswitch | 1.353 | 1.569 | 4.558 | 0.413 |
| tRNA | 0.737 | 0.976 | 4.113 | 0.343 |
| aptamer RNA | 2.171 | 2.533 | 4.997 | 0.480 |
| ribozyme | 1.023 | 1.414 | 4.472 | 0.487 |

Table S14: **Secondary-structure context composition of RNA chains and ligand-binding pockets.** Distribution of DSSR-assigned per-nucleotide secondary-structure contexts in the ligand-interacting RNA chains and in the ligand-binding pockets, for the curated, *non-redundant* HARIBOSS subset (2,908 pockets in 1,620 structures; 2,481,021 chain nucleotides and 31,190 pocket nucleotides). Each nucleotide is assigned to a single mutually exclusive context, so each column sums to 100%. The $\log_2$ enrichment compares each context in pockets relative to the interacting chains.

| **Context** | **RNA chains** | **Pockets** | $\log_2$ **enrichment** |
|---|---|---|---|
| Helix/paired | 55.6 | 47.5 | −0.23 |
| Hairpin loop | 12.3 | 10.0 | −0.30 |
| Internal loop | 14.9 | 14.2 | −0.07 |
| Bulge | 2.1 | 2.3 | +0.15 |
| Junction | 11.8 | 21.0 | +0.83 |
| Single-stranded | 3.3 | 5.0 | +0.59 |

Table S15: **Ligand uniqueness across different interaction environments.** Distribution of unique, shared, and interaction-environment-specific ligands across the raw interaction families identified in HARIBOSS. Interaction families are ordered by decreasing number of unique ligands, and the shared fraction corresponds to the proportion of ligands present in multiple interaction families.

| Interaction family | Total ligands | Shared | Exclusive | Shared fraction |
|---|---|---|---|---|
| `RNA` | 234 | 105 | 129 | 0.45 |
| `ion` | 203 | 103 | 100 | 0.51 |
| `ion`, `protein` | 128 | 81 | 47 | 0.63 |
| `protein` | 114 | 82 | 32 | 0.72 |
| `other` | 56 | 41 | 15 | 0.73 |
| `ion`, `other` | 34 | 34 | 0 | 1.00 |
| `DNA`, `ion`, `protein` | 31 | 20 | 11 | 0.65 |
| `ion`, `other`, `protein` | 29 | 23 | 6 | 0.79 |
| `other`, `protein` | 29 | 24 | 5 | 0.83 |
| `DNA`, `protein` | 12 | 7 | 5 | 0.58 |
| `DNA` | 7 | 3 | 4 | 0.43 |
| `DNA`, `ion`, `other`, `protein` | 6 | 4 | 2 | 0.67 |
| `DNA`, `ion` | 4 | 3 | 1 | 0.75 |
| `DNA`, `other` | 1 | 1 | 0 | 1.00 |
| `DNA`, `other`, `protein` | 1 | 1 | 0 | 1.00 |

Table S16: **Ligand uniqueness across different RNA families.** Distribution of unique, shared, and RNA-family-specific ligands for each RNA family represented in HARIBOSS. The shared fraction corresponds to the proportion of ligands observed in multiple RNA families.

| Type name | Total ligands | Shared | Exclusive | Shared fraction |
|---|---|---|---|---|
| rRNA | 233 | 71 | 162 | 0.305 |
| riboswitch | 101 | 26 | 75 | 0.257 |
| unknown | 86 | 41 | 45 | 0.477 |
| synthetic construct | 74 | 36 | 38 | 0.486 |
| aptamer RNA | 62 | 10 | 52 | 0.161 |
| tRNA | 62 | 48 | 14 | 0.774 |
| mRNA | 38 | 33 | 5 | 0.868 |
| RNA/DNA hybrid | 19 | 13 | 6 | 0.684 |
| TAR RNA | 18 | 11 | 7 | 0.611 |
| ribozyme | 18 | 7 | 11 | 0.389 |
| telomerase RNA | 6 | 2 | 4 | 0.333 |
| SRE RNA | 5 | 4 | 1 | 0.800 |
| snRNA | 3 | 1 | 2 | 0.333 |

Table S17: **Most frequently observed ligands in HARIBOSS.** The ten ligands associated with the largest numbers of RNA-binding pockets in HARIBOSS ($n = 4718$). For each ligand, the table reports the number of associated binding pockets, their proportion of all pockets, and the number of distinct RNA-small molecule complexes in which the ligand is observed.

| Ligand code | Ligand name | # pockets | Percentage | # complexes |
|---|---|---|---|---|
| SPM | Spermine | 957 | 20.3% | 194 |
| PAR | Paromomycin | 416 | 8.8% | 101 |
| GTP | Guanosine-5'-triphosphate | 104 | 2.2% | 79 |
| EPE | HEPES | 99 | 2.1% | 41 |
| LLL | Gentamicin C1A | 98 | 2.1% | 8 |
| GET | Geneticin | 81 | 1.7% | 16 |
| T1C | Tigecycline | 74 | 1.6% | 20 |
| ATP | Adenosine-5'-triphosphate | 70 | 1.5% | 52 |
| SAM | S-Adenosylmethionine | 66 | 1.4% | 45 |
| SAH | S-Adenosyl-L-Homocysteine | 63 | 1.3 % | 35 |

### References


(1) Naineni, S. K.; Liang, J.; Hull, K.; Cencic, R.; Zhu, M.; Northcote, P.; Teesdale-Spittle, P.; Romo, D.; Nagar, B.; Pelletier, J. Functional mimicry revealed by the crystal structure of an eIF4A:RNA complex bound to the interfacial inhibitor, desmethyl pateamine A. *Cell Chemical Biology* **2021**, *28*, 825–834.e6.

(2) Temiakov, D.; Patlan, V.; Anikin, M.; McAllister, W. T.; Yokoyama, S.; Vassylyev, D. G. Structural Basis for Substrate Selection by T7 RNA Polymerase. *Cell* **2004**, *116*, 381–391.

(3) Freisz, S.; Lang, K.; Micura, R.; Dumas, P.; Ennifar, E. Binding of Aminoglycoside Antibiotics to the Duplex Form of the HIV-1 Genomic RNA Dimerization Initiation Site. *Angewandte Chemie International Edition* **2008**, *47*, 4110–4113.

(4) Lipinski, C. A.; Lombardo, F.; Dominy, B. W.; Feeney, P. J. Experimental and computational approaches to estimate solubility and permeability in drug discovery and development settings. *Advanced Drug Delivery Reviews* **1997**, *23*, 3–25.

(5) Veber, D. F.; Johnson, S. R.; Cheng, H.-Y.; Smith, B. R.; Ward, K. W.; Kopple, K. D. Molecular Properties That Influence the Oral Bioavailability of Drug Candidates. *Journal of Medicinal Chemistry* **2002**, *45*, 2615–2623.

(6) Chen, L.-L.; Kim, V. N. Small and long non-coding RNAs: Past, present, and future. *Cell* **2024**, *187*, 6451–6485.

(7) Hughes, J. D. et al. Physiochemical drug properties associated with in vivo toxicological outcomes. *Bioorganic & Medicinal Chemistry Letters* **2008**, *18*, 4872–4875.
